# Three-dimensional microenvironment regulates gene expression, function, and tight junction dynamics of iPSC-derived blood-brain barrier microvessels

**DOI:** 10.1101/2021.08.27.457975

**Authors:** Raleigh M. Linville, Matthew B. Sklar, Gabrielle N. Grifno, Renée F. Nerenberg, Justin Zhou, Robert Ye, Jackson G. DeStefano, Zhaobin Guo, Ria Jha, John J. Jamieson, Nan Zhao, Peter C. Searson

## Abstract

The blood-brain barrier (BBB) plays a pivotal role in brain health and disease. In the BBB, brain microvascular endothelial cells (BMECs) are connected by tight junctions which regulate paracellular transport, and express specialized transporter systems which regulate transcellular transport. However, existing *in vitro* models of the BBB display variable physiological accuracy across a wide range of characteristics including gene/protein expression and barrier function. Here, we use an isogenic family of fluorescently-labeled iPSC-derived BMEC-like cells (iBMECs) and brain pericyte-like cells (iPCs) within two-dimensional confluent monolayers (2D) and three-dimensional (3D) tissue-engineered microvessels to explore how 3D microenvironment regulates gene expression and function of the *in vitro* BBB. We show that 3D microenvironment (shear stress, cell-ECM interactions, and cylindrical geometry) increases BBB phenotype and endothelial identity, and alters angiogenic and cytokine responses in synergy with pericyte co-culture. Tissue-engineered microvessels incorporating junction-labeled iBMECs enable study of the real-time dynamics of tight junctions during homeostasis and in response to physical and chemical perturbations.

## Introduction

Brain microvascular endothelial cells (BMECs), along with zonation-specific supporting cells, constitute the blood-brain barrier (BBB) and regulate transport into and out of the brain. The highly specialized BMECs possess regulate transport via expression of: (1) tight junctions (TJs) which block paracellular transport (e.g. claudin-5, occludin, ZO1), (2) efflux pumps which limit passive transcellular transport (e.g. P-gp, BCRP), (3) transporter systems which regulate transcellular nutrient transport (e.g. GLUT-1, Lat-1), and (4) specialized fatty acid transporters which restrict vesicular transcytosis (e.g. Mfsd2a) [1]. Strategies to bypass the BBB for drug delivery include transient tight junction disruption by hyperosmotic agents or focused ultrasound, efflux pump inhibition to increase substrate permeability, and trojan-horse approaches that hijack receptor-mediated transporter systems [2]. While transient loss of barrier function may facilitate treatment of diseased neurons or cancerous cells, permanent changes in barrier function can occur during neurological disease resulting in positive feedback to disease progression [3].

While animal models are indispensable for studies of the BBB, there are considerable species-to-species variations and technical limitations [4, 5], which can be at least partially addressed using *in vitro* BBB models. However, the lack of appropriate cell sources remains a major obstacle to their development [6-8]. Primary and immortalized BMECs have established endothelial origin, but physiological barrier function is rarely achieved [1, 9]. Similarly, transendothelial electrical resistance (TEER), a measure of barrier integrity, is expected to be 1,500 - 8,000 Ω cm^2^ *in vivo* [10, 11], but is usually less than 200 Ω cm^2^ for primary and immortalized BMECs [12-15]. Primary and immortalized cell sources have other disadvantages including batch-to-batch variability [12], loss of phenotype during *in vitro* culture [16], and complicated isolation procedures which limit scalability [4]. To address these limitations, a diverse array of differentiation schemes have emerged to generate BMEC-like cells from induced pluripotent stem cells (iBMECs) (summarized in [17]). These protocols rely on endothelial/mesodermal specification followed by brain endothelial specification, achieved via a combination of Wnt/β-catenin, VEGF, and retinoic acid-induced signaling [18-22]. While recent evidence suggests that iBMECs possess a component of epithelial identity [23], these cells remain a critical source for BBB modeling as they exhibit functional barrier characteristics including high TEER, low permeability, efflux activity, and nutrient transport, while enabling highly scalable and patient-specific experimentation [1, 17]. Recent approaches to drive brain endothelial identity by transcription factor (TF) reprogramming of iPSC-derived cells [23, 24] or by chemical exposure to Wnt agonists/ligands or TGF-beta inhibitors [25, 26] hold promise, but have been unable to achieve physiological barrier function with TEER values typically < 200 Ω cm^2^.

*In vitro* BBB models have largely relied on two-dimensional (2D) confluent monolayers which lack many microenvironmental cues present within the cerebrovasculature. While the roles of shear stress [27-29] and co-cultured pericytes [30, 31] have been explored in microfluidic chip BBB models, the interplay of other microenvironmental cues present *in vivo* (cell-ECM interactions, transmural pressure, and cylindrical geometry) is not well established. Here, we differentiated iBMECs from an isogenic family of fluorescently-labeled iPSCs to enable visualization of tight junctions, cytoskeleton, or cell membranes using live-cell imaging in a three-dimensional (3D) tissue-engineered BBB model. The two major objectives were: (1) to characterize how 3D microenvironment augments iBMEC phenotype, and (2) to use this platform to visualize the dynamics of tight junctions during homeostasis and injury. We find that the 3D microenvironment improves the stability and strength of iBMEC barrier properties, increases expression of endothelial transcripts, and improves correlation to human BMEC and endothelial cell transcriptomes. We address conflicting reports of angiogenic and cytokine responses in iBMECs (in synergy with pericyte co-culture), and show that the 3D microenvironment imparts unique functional responses. To highlight the applications of iBMEC models, we mapped the dynamics of tight junctions during homeostasis, physical injury, and chemical injury. The responses of iBMEC monolayers were found to be unique in 3D microenvironment and across different perturbations (ablation, oxidative stress, and peptide exposure).

## Results

### Differentiation and characterization of fluorescently-labeled isogenic iBMECs

To enable live-cell imaging studies of the BBB during homeostasis and in response to physical, chemical, and cellular perturbations, we developed an isogenic family of BMEC-like cells from CRISPR-edited source iPSCs (iBMECs). We reproduced iBMEC differentiation protocols [19, 22] across three isogenic cell lines with fluorescently-labeled ZO1 (iBMEC-TJs), plasma membrane (iBMEC-PMs), and β-actin (iBMEC-ACTBs) (**Fig. 1A**). Following differentiation, iBMECs were sub-cultured on collagen IV and fibronectin-coated plates where we observed cobblestone morphology and the formation of confluent monolayers enabling real-time imaging of tight junctions, plasma membrane, and actin cytoskeleton (**Fig. S1A**). iBMECs were incorporated within 2D and 3D models to compare barrier properties, gene expression, and functional responses (**Figs. 1-3**), and to visualize the dynamics of tight junctions during homeostasis and injury (**Figs. 4-6**). To provide an overview of barrier function we selected representative substrates for active transport (glucose/GLUT1), efflux pumps (rhodamine 123/P-gp), and paracellular transport (Lucifer yellow and dextran) (**Figs. S1 - S3**).

**Figure 1.**
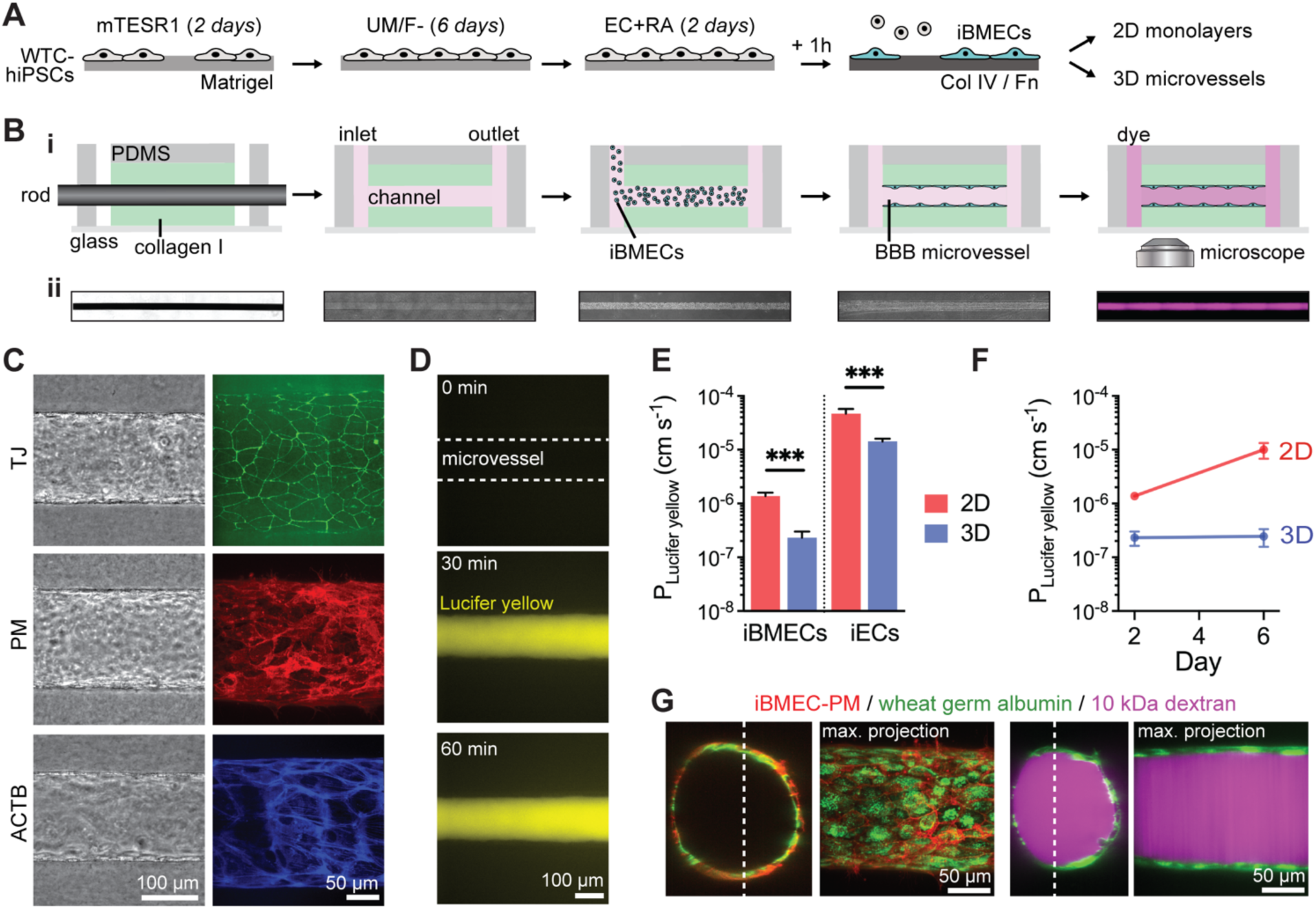
Three-dimensional BBB model incorporating iBMECs displays stable and low paracellular permeability. (A) Schematic illustration of the differentiation protocol. WTC hiPSCs were sequentially treated with mTESR1 media (2 days), unconditioned media without bFGF (UM/F-; 6 days) and retinoic acid-supplemented endothelial media (EC+RA; 2 days). iBMECs were purified by a one-hour sub-culture on collagen IV and fibronectin-coated plates, and then used within 2D or 3D models. (B) Fabrication and imaging of iBMEC microvessels. 150 μm diameter channels patterned in collagen I were seeded with iBMECs to form a confluent endothelium. Insets below show phase contrast and fluorescence images along the experimental timeline. (C) Representative images show organization of iBMEC microvessels with localized tight junctions (TJ), plasma membrane (PM), and actin cytoskeleton (ACTB) expression under confocal imaging (max. intensity projection shown). (D) Representative images following perfusion with Lucifer yellow. (E) Permeability of Lucifer yellow on day 2 across microvessel conditions: iBMEC microvessels (*n* = 14), iBMEC Transwells (*n* = 11), isogenic iEC microvessels (*n* = 4 microvessels), and iEC Transwells (*n* = 4). (F) Time course of Lucifer yellow permeability over one week for iBMEC microvessels (*n* = 14 and 4, respectively) and iBMEC Transwells (*n* = 11). (G) Confocal images of iBMEC-PM microvessels highlighting the glycocalyx and showing solute perfusion in the lumen. (Left) Wheat germ albumin staining of iBMEC glycocalyx. (Right) 10 kDa dextran not observed to accumulate within glycocalyx or iBMECs (60 minutes after exposure). Data are presented as mean ± SEM. *** *p* < 0.001. See also **Figure S1-3**; permeability experiments were conducted across TJ, PM, and ACTB-labeled source cells.

### Barrier function in 2D Transwells

Motivated by a desire to minimize reagent use while maintaining cellular fidelity, we differentiated iBMECs in parallel using 1 mL or 2 mL of medium to determine impact of media volume on cell phenotype (Fig. S1B). With reduced media volume during differentiation, iBMECs displayed improved phenotype as measured by transendothelial electrical resistance (TEER), paracellular and transcellular permeability, protein expression, and gene expression (see *Note S1* and **Fig. S1**). Mean TEER values for all cell lines and differentiations were within the range of *in vivo* measurements in animal models and theoretical calculations (1,500 - 8,000 Ω cm^2^) [1]. Regardless of media volume, TEER values decreased over six days, matching previous observations [19, 21, 22] and suggesting limited phenotypic stability on Transwells.

Reduced media volume lowered Lucifer yellow (444 Da) permeability measured two days after culturing iBMECs onto Transwells (p = 0.043) (**Fig. S1F**), while not substantially altering the permeability of 10 kDa dextran (p = 0.288) (**Fig. S1G**). Comparison of the two media volumes revealed no difference in the ratio of apical-to-basolateral permeability for glucose transport (p = 0.353) and no difference in basolateral-to-apical permeability for rhodamine 123 (p = 0.054), indicating no difference in GLUT1 and P-gp efflux activity (**Fig. S1H**). Gene and protein expression of endothelial and BMEC markers were similar regardless of media volume (**Fig. S1I-J**). Together these results show that reduced media volume can be used to differentiate iBMECs that display functional hallmarks of human BMECs. We also showed that media volume effects are maintained following cryopreservation, using an independent serum lot, and using an independent iPSC source, and explored the relationship between TEER and permeability across our datasets (see *Note S2* and **Fig. S2**). Additionally, the mechanistic underpinnings of the reduced media volume effect remain to be fully elucidated, but are likely related to soluble factors secreted during the initial UM/F-phase of differentiation (see *Note S3* and **Fig. S3**).

### Barrier function in 3D iBMEC microvessels

Tissue-engineered microvessels were generated by seeding iBMECs into 150 μm diameter channels patterned in collagen I (**Fig. 1B-i**). iBMECs formed confluent ∼1 cm-long microvessels free from defects as evidenced by a lack of 10 kDa dextran leakage (**Fig. 1B-ii**). The library of fluorescently-labeled isogenic iPSCs enabled real-time imaging of tight junctions, plasma membrane, and actin cytoskeleton via epifluorescence or confocal microscopy (**Fig. 1C**). iBMEC microvessels maintained low Lucifer yellow (444 Da) permeability (∼2.3 × 10^−7^ cm s^-1^) independent of iPSC source, with no visible sites of paracellular leakage indicating formation of a confluent monolayer connected by tight junctions (**Fig. 1D**). Compared to induced isogenic endothelial cell (iEC) microvessels, iBMEC microvessels displayed ∼60-fold lower permeability demonstrating acquisition of brain-specific barrier properties (**Fig. 1E**). For both iBMECs and iECs, 3D culture was associated with reduced paracellular permeability (*p* < 0.001) (**Fig. 1E**). The permeability of Lucifer yellow across 2D iBMEC monolayers was significantly higher than in 3D microvessels on day 2, and increased ∼6-fold at day 6 (*p* = 0.078) (**Fig. 1F, S2C**), matching observations of declining TEER values (**Fig. S1A, S2A**). In contrast, 3D iBMEC microvessels maintained stable barrier function in the absence of co-cultured cells over at least six days (*p* > 0.924) (**Fig. 1F**). The lower permeability in 3D may be due to a number of factors, including shear stress, substrate stiffness, curvature, or the perimeter effect (see *Note S4*) [32]. The incorporation of iBMEC-PMs into iBMEC microvessels enabled identification of the luminally localized glycocalyx and exclusion of 10 kDa dextran from the endothelium (**Fig. 1G**). These results suggest that the 3D microenvironment promotes physiological and stable paracellular barrier function.

### Microenvironmental regulation of gene expression

The effects of the 3D microenvironment on iBMEC gene expression were evaluated by comparison of bulk RNA sequencing of iBMECs in 3D microvessels and 2D monolayers, and distinguished from the effects of shear stress alone by comparison to previously published RNA transcriptomes in a 2D microfluidic chip model [28] (**Fig. 2A**). Bulk RNA sequencing of iBMECs across two independent iPSC sources in 3D microvessels revealed 497 upregulated genes and 228 downregulated genes compared to iBMECs in 2D static monolayers (ρ = 0.975) (**Fig. 2B, Data S1**). Upregulated transcripts included canonical endothelial genes (*TIE1, FLI1, ICAM1, SERPINE1*), transcription factors (*GLIS3, AFF3*), metallothioneins (*MT2A, MT1E*), mediators of the canonical Wnt/beta-catenin signaling pathway (*WNT7A, WNT2B*), cytokines and cytokine receptors (*CX3CL1, CXCL16, IL23A, IL32, IL6R, IL20RB, IL15RA*), monooxygenases involved in drug metabolisms (*CYP1A1, CYP1B1*), integrins (*ITGB6, ITGA2, ITGBL1*), and transforming growth factor-beta (TGFB) family members (*TGFA, TGFB1, TGFB2*); downregulated transcripts included retinoid X receptors (*RXRA*), retinoic acid receptors (*RARG*), and collagen family members (*COL2A1, COL26A1*), among others.

**Figure 2.**
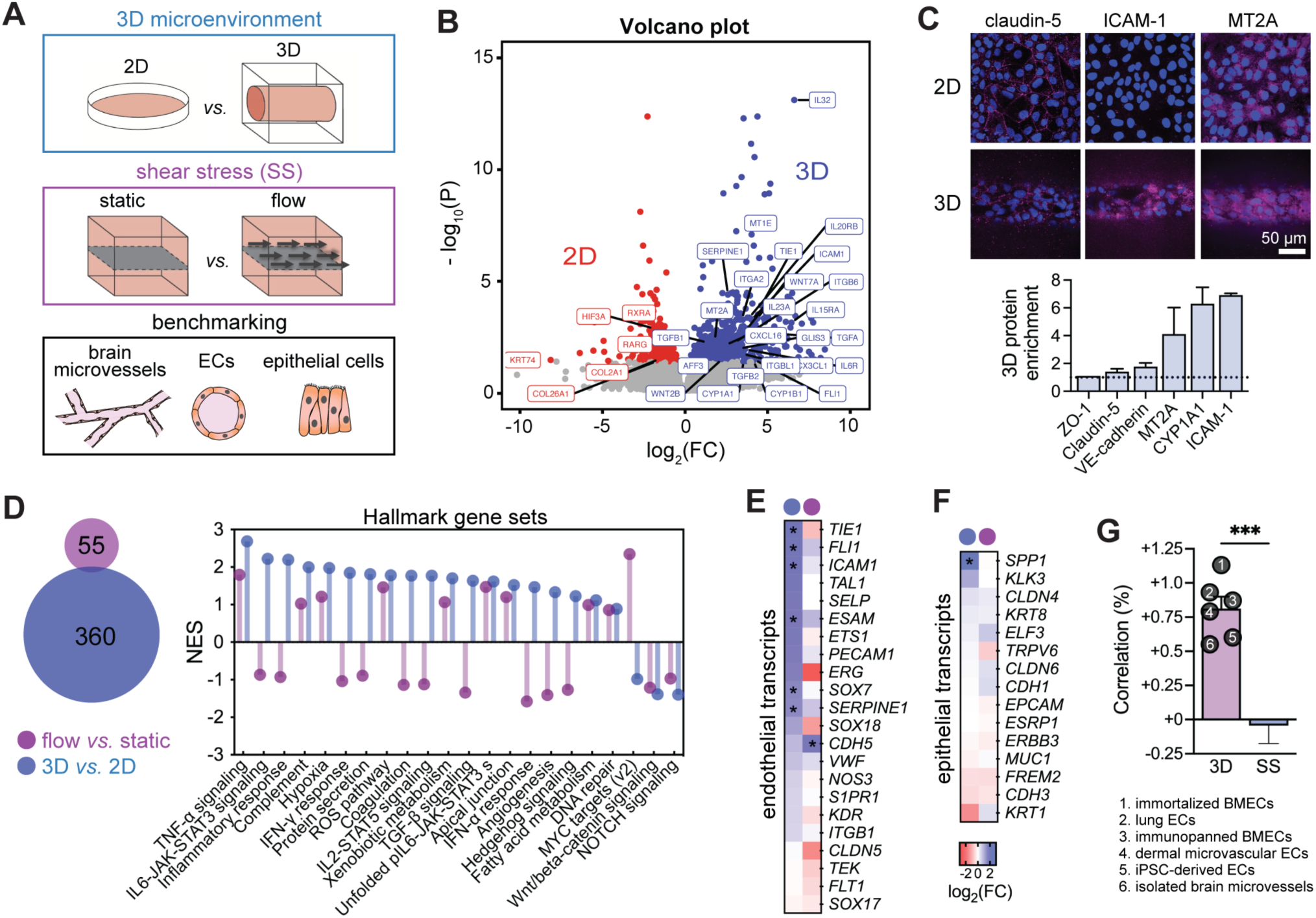
Three-dimensional microenvironment regulates iBMEC gene expression. (A) To elucidate the role of 3D microenvironment on iBMEC gene expression, 2D and 3D models were compared by paired bulk RNA sequencing. These findings are benchmarked to previous work on the role of shear stress (SS) [29] and to datasets for brain microvessels, endothelial cells, and epithelial cells. (B) Volcano plots depicting significantly (adjusted *p* < 0.05, Wald test with Benjamini-Hochberg correction) upregulated genes (blue) and downregulated genes (red) in iBMECs in 3D microvessels versus 2D monolayers (*n* = 5 per condition). The five paired 2D/3D replicates encompass three WTC iBMECs (one for each WTC iPSC source) and two BC1 iBMECs (an independent iPSC source). Selected genes are labeled. (C) Semi-quantitative validation of protein enrichment in 3D microvessels. Representative immunofluorescence images for claudin-5, ICAM-1, and MT2A in 2D and 3D are shown. Data is normalized to nuclei signal and endothelium area and presented as mean ± SEM (*n* = 2). (D) Comparison of upregulated genes between the 3D microenvironment and 2D monolayers under shear stress. The Venn diagram shows overlap of genes upregulated by the 3D microenvironment or shear stress by log_2_FC > 1. The lollipop plot highlights normalized enrichment scores (NES) of select Hallmark gene sets by 3D microenvironment and shear stress (see Fig. S4A for all 50 gene sets). (E-F) Heatmaps of changes in endothelial and epithelial transcript abundance due to 3D microenvironment versus shear stress; DEGs are labeled with an asterisk. (G) iBMECs were benchmarked to datasets for brain microvessels, endothelial cells, and epithelial cells (see Fig. S5 for complete analysis). 3D microenvironment increases the correlation to brain microvessels and endothelial cells, while shear stress (SS) alone does not. Data are presented as mean ± SEM. *** *p* < 0.001. See also **Figure S4-6** and **Data S1**.

We validated upregulation of several genes at the protein level using immunocytochemistry; this approach provides critical information on protein localization, and can be conducted on models comprised of small numbers of cells, where Western blot is technically burdensome. Metallothionein-2 (*MT2A*), a cytochrome P450 super family member (*CYP1A1*), and ICAM-1 (*ICAM1*) were confirmed to be enriched 4- to 6-fold in 3D microvessels based on semi-quantitative analysis of relative protein expression normalized to the fluorescence of DAPI-stained nuclei (**Fig. 2C**). *CYP1A1* and *MT2A* expression had previously been found to be shear stress dependent in primary or iPSC-derived BMECs [28, 33-35]. We confirmed that the tight junction proteins claudin-5 (*CLDN5*) and zona occludens-1 (*TJP1*), which were not substantially upregulated at the gene level by the 3D microenvironment, were also similarly expressed at the protein level (**Fig. 2C**).

In contrast to studies of the role of laminar flow on 2D monolayers [27, 28], tissue-engineered iBMEC microvessels were exposed to stimuli including cell-ECM interactions, cylindrical geometry, and transmural pressure. We identified 360 genes upregulated by greater than two-fold compared to only 55 genes upregulated to this same level by 2.4 dyne cm^-2^ shear stress in a previous study [28]; less than 1% of upregulated genes were shared, indicating unique modes of gene expression changes (**Fig. 2D**). To predict functional differences induced by the 3D microenvironment and shear stress, we conducted gene set enrichment analysis (GSEA) of hallmark gene sets from the Molecular Signatures Database (MSigDB) [36] (**Fig. 2D, S4A)**. Many hallmark gene sets were oppositely enriched by 3D microenvironment compared to shear stress, including IL6-JAK-STAT3 signaling, inflammatory response, IFN-γ response, protein secretion, coagulation, IL2-STAT5 signaling, TGF-β signaling, IFN-α response, angiogenesis, and hedgehog signaling, further supporting distinct functional changes due to the 3D microenvironment. Gene sets enriched by both 3D microenvironment and shear stress included TNF-α signaling, complement, hypoxia, ROS pathway, and xenobiotic metabolism.

Given recent findings on the partial epithelial identity of iBMECs [37], we explored changes in endothelial and epithelial gene expression induced by the 3D microenvironment and shear stress. Endothelial and epithelial gene lists were curated from the literature [37, 38]. Endothelial transcripts (which have previously been reported to be attenuated in iBMECs compared to other endothelial cell sources [37, 39]) were broadly increased by the 3D microenvironment (**Fig. 2E**). These transcripts displayed on average 2-fold enrichment by the 3D microenvironment, while no enrichment was observed by shear stress alone (*p* < 0.0001) (**Fig. 2E, S4B**) [28]. The 3D microenvironment and shear stress did not dramatically alter expression of epithelial genes (*p* = 0.699) (**Fig. 2F, S4B**); demonstrating that endothelial identity can be independently augmented from epithelial identity.

### Benchmarking gene expression of 3D iBMEC microvessels

To facilitate comparison of iBMECs to endothelial and epithelial cell sources, we conducted meta-analysis of global gene expression collected across bulk RNA-sequencing studies (**Fig. 2A, S5**). We performed in-house sequencing of primary human dermal microvascular endothelial cells (HDMECs) and iPSC-derived endothelial cells (iECs) generated by transient expression of *ETV2* after mesodermal induction [40]. Previously published RNA transcriptomes were also obtained from the NCBI Gene Expression Omnibus (GEO) from the following studies: (1) immortalized human BMECs [41], (2) primary human lung ECs [42], (3) immunopanned human BMECs [43], (4) human brain microvessels [4], (5) primary human bronchial epithelial cells [44], and (6) the caco-2 epithelial cell line [45] (**Table S1**). Spearman correlation of variance stabilized transcript expression identified similarities between immunopanned human BMECs and human brain microvessel gene profiles, with separate clusters of iBMECs, endothelial cells, and epithelial cells (**Fig. S5A**). iBMECs clustered more closely with epithelial transcriptomes as previously reported [23]. However, iBMECs, endothelial cells, and epithelial cells displayed unique gene expression profiles compared to human microvessel samples with Spearman correlation coefficients (ρ) ranging from 0.726 to 0.817. Endothelial cell types including immortalized BMECs, iECs, HDMECs, and lung ECs were highly similar to each other (ρ > 0.9); however, none of these cell sources exhibit TEER values resembling *in vivo* measurements [1]. In contrast, iBMECs had unique gene expression profiles compared to the group of endothelial cells suggesting a reduction in endothelial identity [23], but exhibited physiologically high TEER.

We compared changes in the correlation to datasets for endothelial and epithelial cells between 3D microenvironment and shear stress (**Fig. 2G, S5A-B**). The 3D microenvironment increased Spearman correlation coefficients to endothelial datasets by ∼0.75%, significantly more than shear stress alone (*p* = 0.0003), suggesting that other components of the 3D microenvironment may be important in promoting brain endothelial identity [28]. Correlation to epithelial datasets was not strongly affected by either 3D microenvironment or shear stress (**Fig. S5B**). To characterize the magnitude of gene expression changes induced by 3D microenvironment, we compared Spearman correlation coefficients across recent studies of shear stress, chemical induction, and TF reprogramming [23, 25, 26, 28] (**Fig. S5C-D**). The magnitude of gene expression changes induced by the 3D microenvironment (ρ = 0.984 between 3D and 2D iBMECs) was similar to other perturbations explored to induce brain endothelial identity, including shear stress (ρ = 0.980 between 2.4 dyne cm^-2^ and static iBMEC monolayers) [28], agonism of Wnt/β-catenin signaling in endothelial progenitor cells (ρ = 0.977 between EPCs exposed to CHIR99021 or DMSO) [26], and inhibition of TGFβ receptor in iECs (ρ = 0.983 between iECs exposed to RepSox or DMSO) [25] (**Fig. S5D**). However, TF reprogramming of iBMECs using ETS transcription factors [23] induced a stronger change in gene expression (ρ = 0.847 compared to 2D iBMEC monolayers), with average > 5 log_2_FC enrichment of endothelial transcripts and downregulation of epithelial transcripts (**Fig. S5E**). Abundance measurements of endothelial and epithelial transcripts across cell sources and model types (see **Figure S6)** provide further support that the 3D microenvironment increases endothelial transcripts in iBMECs compared to 2D and epithelial cells, but at lower levels than other endothelial cell sources.

### Microenvironmental regulation of angiogenic and cytokine response

To explore the synergy between microenvironmental factors regulating iBMEC phenotype, we incorporated iPSC-derived pericytes into our tissue-engineered BBB model. Pericytes exert diverse influences on BMECs, including regulation of angiogenesis and leukocyte infiltration [46-48]. An isogenic model was generated by seeding neural crest iPSC-derived pericytes (iPCs) (generated using published protocols [49]) into microvessels for one day prior to seeding iBMECs. The iPCs maintained direct cell-cell contact with iBMECs and were located along the abluminal surface of the endothelium (**Fig. 3A**). iPCs expressed NG2 and PDGFRβ at the gene and protein level (**Fig. S4C, D**). Given that the TNF-α signaling and angiogenesis hallmark gene sets were upregulated by the 3D microenvironment, we explored the response to angiogenic factors and cytokines by perfusing microvessels with 20 ng mL^-1^ bFGF for 48 h or 5 ng mL^-1^ TNFα for 24 h (**Fig. 3B**).

**Figure 3.**
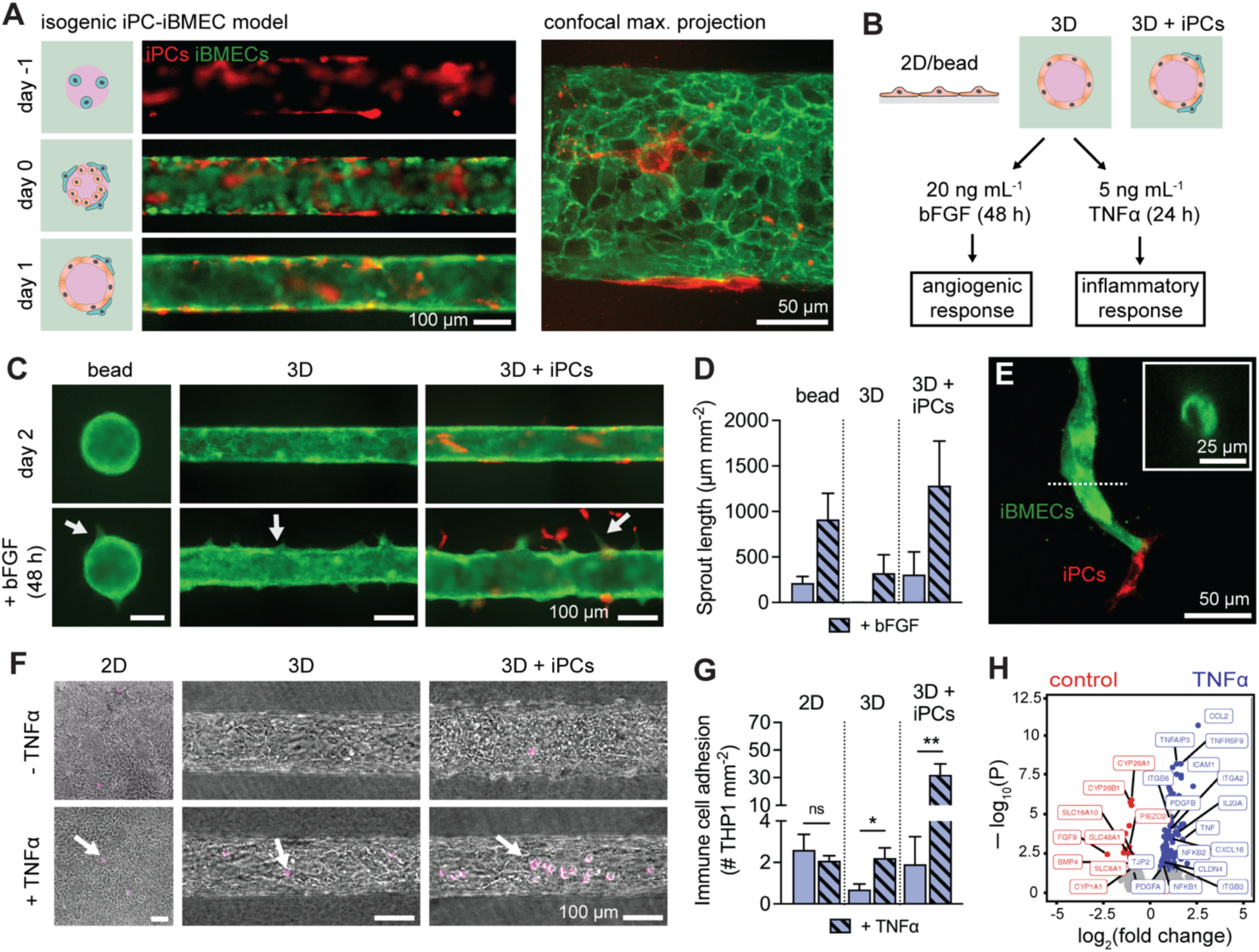
Three-dimensional microenvironment and pericyte co-culture synergistically alter angiogenic and cytokine responses. (A) The combination of ACTB-labeled iBMECs and PM-labeled iPCs enables creation of an isogenic co-culture BBB model. (i) The model was formed via sequential seeding of pericytes and then endothelial cells, to achieve a 1:3 ratio. (ii) Pericytes localized abluminal to iBMEC-PM. (B) Experimental design: 2D, 3D and 3D models co-cultured with pericytes were exposed to bFGF or TNFα to quantify cytokine and angiogenic responses, respectively. (C) Representative images of iBMEC response to bFGF. Angiogenic sprouts appear over two days of exposure (white arrows). (D) Quantification of angiogenic sprout length following 48 h exposure to 20 ng mL^-1^ bFGF. Data collected across bead assay (*n* = 4), 3D microvessels (*n* = 4), and pericyte co-cultured 3D microvessels (*n* = 3). (E) Pericytes located at the leading edge of an angiogenic sprout. Inset: representative lumen cross-section at location of dotted line. (F) Representative images of iBMEC response to TNFα. Experiments were performed on BC1 iBMECs paired with RFP-tagged WTC iPCs to enable detection of adherent immune cells (white arrows). (G) Quantification of immune cell adhesion following 24 h exposure to 5 ng mL^-1^ TNFα. Data collected across 2D monolayers (*n* = 8), 3D microvessels (*n* = 5), and pericyte co-cultured 3D microvessels (*n* = 4). (H) Volcano plots showing significantly (adjusted *p* < 0.05, Wald test with Benjamini-Hochberg correction) upregulated genes (blue) and downregulated genes (red) following 24 h exposure to 5 ng mL^-1^ TNFα in 3D microvessels (*n* = 2 biological replicates). Selected genes are labeled. Data are presented as mean ± SEM. * *p* < 0.05, ** *p* < 0.01, *** *p* < 0.001, and **** *p* < 0.0001. See also **Figure S7** and **Data S1**.

The angiogenic response was compared across three conditions (microvessels, microvessels co-cultured with pericytes, and a bead assay). As previously observed, iBMECs were responsive to growth factors, including bFGF [50] (**Fig. 3C**). In the absence of bFGF, iBMEC microvessels displayed no sprouting, while bFGF treatment resulted in the formation of sprouts with a visible lumen (**Fig. 3D**). iBMEC/PC microvessels displayed elevated responsiveness to bFGF with longer sprouts, visible lumens, and pericytes located at the leading edge (**Fig. 3E**).

Previous studies of iBMECs have reported conflicting degrees of responsiveness to inflammatory cytokines [21, 23, 51, 52]. Here, we assessed the response of 2D iBMEC monolayers, iBMEC microvessels, and iBMEC/iPC microvessels to tumor necrosis factor-alpha (TNFα) by measuring the adherence of monocyte-like cells (THP1s) (**Fig. 3F**). In 2D monolayers, the density of adhered THP1s was high and did not increase following TNFα exposure (*p* = 0.5136). In 3D, immune cell adhesion was low under control conditions, but increased ∼3-fold following TNFα exposure (*p* = 0.0259) (**Fig. 3G**). Furthermore, in the presence of pericyte co-culture, immune cell adhesion was dramatically enhanced upon TNFα exposure (∼16-fold) (*p* = 0.0094). These results suggest that the dynamics of immune cell adhesion are synergistically modulated by the 3D microenvironment and pericytes.

We also compared gene expression of iBMEC microvessels with and without TNFα exposure by bulk RNA sequencing, identifying 136 upregulated genes and 35 downregulated genes (ρ = 0.971) (**Fig. 3H, Data S1**). Upregulated genes included NF-κB family members (*NFKB1, NFKB2*), tumor necrosis factor family members (*TNF, TNFAIP3, TNFRSF9*), endothelial transmembrane proteins facilitating leukocyte transmigration (*ICAM1*), cytokines (*CCL2, CXCL16, IL23A*), growth factors (*PDGFA, PDGFB*), cell-cell junctions (*CLDN4, TJP2*) and integrins (*ITGB3, ITGB6, ITGA2*). Downregulated genes included solute carrier (SLC) membrane transporters (*SLC8A1, SLC48A1, SLC16A10*), monooxygenases involved in drug metabolisms (*CYP1A1, CYP26B1, CYP26A1*), mechanosensitive ion channels (*TRPV6, PIEZO2*), and growth factors (*BMP4, FGF9*).

### Dynamics of iBMEC monolayers in tissue-engineered microvessels

To study the dynamic response of iBMECs to different physiologically-relevant perturbations, we performed live-cell imaging of iBMEC-TJs in tissue-engineered microvessels. ZO1 (*TJP1*) is a scaffolding protein that connects transmembrane TJ proteins to the actin cytoskeleton [53], enabling measurements of morphology, motility, and turnover (**Fig. 4A**). The dynamic response of these metrics integrates various biological pathways including wound healing and stress responses. Fluorescence imaging revealed that during homeostasis tight junctions were dynamic, with small fluctuations in the position of cell-cell junctions between adjacent cells; most cells were quiescent (∼97%) during imaging and displayed highly stable morphology (**Fig. 4B**).

**Figure 4.**
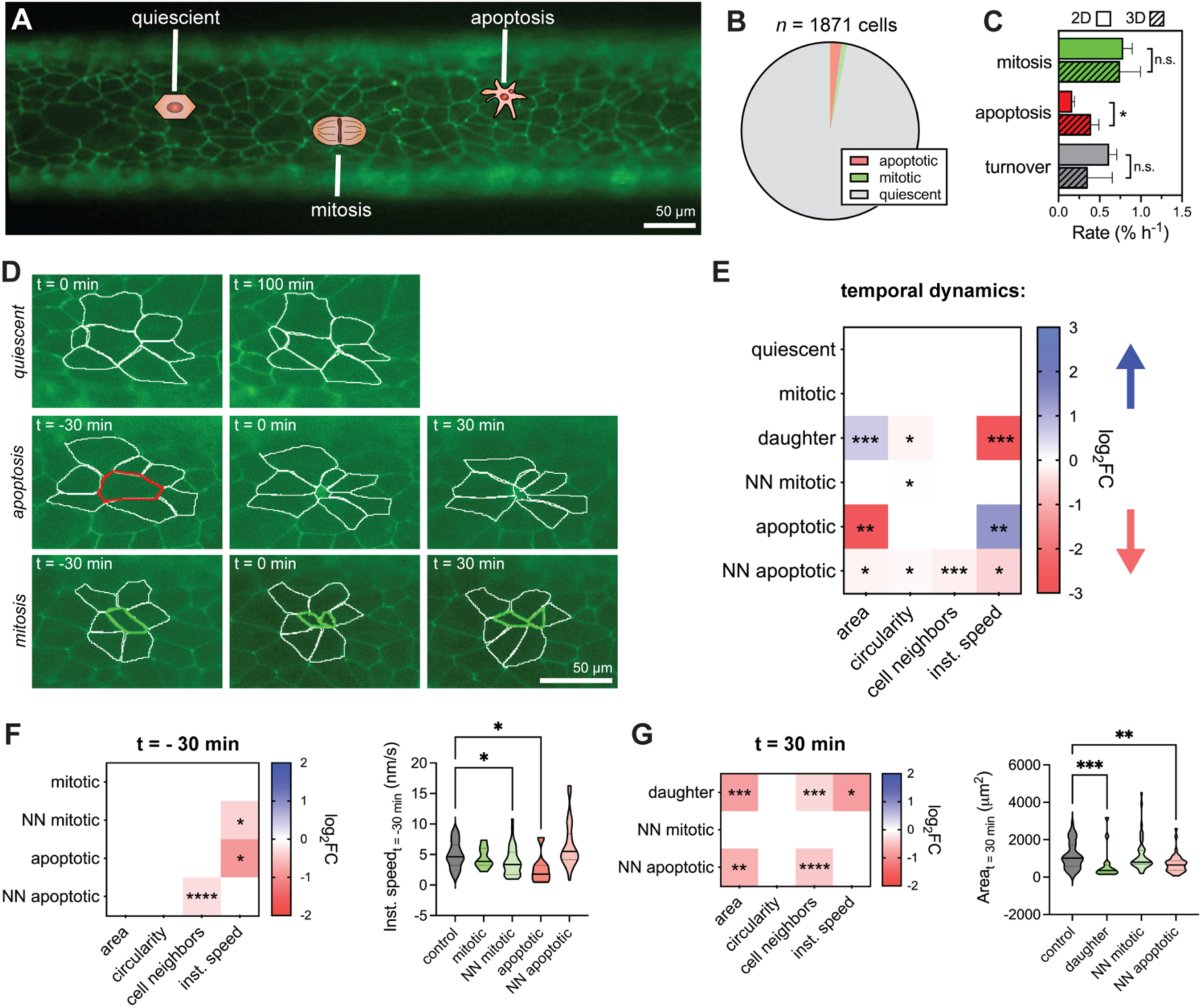
Monolayer dynamics in three-dimensional iBMEC microvessels. (A) Tight junction (TJ)-labeled iBMECs support tracking of monolayer dynamics in real time. (B) Apoptotic, mitotic, and quiescent cell populations identified from imaging of *n* = 1871 cells from *n* = 6 iBMEC-TJ microvessels. (C) Comparison of mitosis, apoptosis, and turnover rates between iBMECs in 3D microvessels (*n* = 7) and 2D monolayers (*n* = 6). Significance calculated using a Mann-Whitney test. (D) Apoptotic, mitotic, and quiescent cells are identified from changes in tight junction fluorescence and phase contrast imaging over 100 minutes for quiescent cells and 60 minutes for apoptotic and mitotic cells, with an interval of 5 minutes. (E) Heatmap summarizing morphological changes during imaging. Only squares corresponding to a statistically significance change are shaded by magnitude of change (log_2_FC) and direction of change (blue: increasing over time, red: decreasing over time) labeled with asterisks representing significance as calculated using the maximally significant result between a Wilcoxon matched-pairs signed rank test to compare first and last time points and using an F test to determine if linear regression of the morphological time course displayed a statistically non-zero slope. (F, G) Heatmaps comparing morphological metrics of cell types before (t = - 30 min) and after (t = 30 min) cellular events. Only squares corresponding to a statistically significance comparison are shaded by magnitude of change (log_2_FC) and direction of change (blue: higher than quiescent cells, red: lower than quiescent cells) labeled with asterisks representing significance calculated using a Kruskal-Wallis test and post hoc Dunn’s multiple comparisons test relative to quiescent cells. Specific violin plots of interest are also shown. Data are presented as mean ± SEM. NN – nearest neighbor. * *p* < 0.05, ** *p* < 0.01, *** p < 0.001, and **** p < 0.0001. See also **Figure S8-10** and **Data S2**; all analysis conducted on TJ-labeled iBMECs.

Apoptotic cells were visible based on nuclear fragmentation and formation of apoptotic bodies under phase contrast imaging, and subsequent cell collapse under fluorescence imaging (t = 0 defined as end of cell collapse) (**Fig. S8A**). Mitotic cells were visible based on chromosomal alignment and separation under phase contrast imaging, and cytokinesis under fluorescence imaging (t = 0 defined as separation of daughter cells via tight junction) (**Fig. S8B**). In homeostasis, the net monolayer turnover was very low (typically < 0.5 % h^-1^). Interestingly, while rates of mitosis and turnover were similar between 2D and 3D culture (*p* > 0.05), apoptosis was higher in the 3D microenvironment (*p* = 0.0140) (**Fig. 4C, S8A-B**), possibly related to enrichment of apoptosis-related genes from GSEA. In addition, imaging during permeability measurements showed that iBMECs were able to reorganize their junctions to maintain barrier function during apoptosis and mitosis, thereby preventing transient leakage.

We recorded the morphology of 7 quiescent, 8 mitotic, and 7 apoptotic cells along with all their nearest neighbors (NN) and progeny (*n* = 151 cells) with 5-minute resolution for 30 minutes before and after these events (*t* = 0 defined above) (**Fig. 4D, S9, Data S2**). From principal component analysis (PCA), we found that monolayer dynamics were dependent on four key parameters: (1) cell size (area/perimeter), (2) cell shape (circularity/aspect ratio), (3) cell motility (instantaneous speed), and (4) number of cell neighbors (local density of cells in a monolayer) (**Fig. S8C**). Although the number of cell neighbors was highly correlated with cell area, there were distinct contributions of cell neighbors to single cell morphology warranting inclusion as a separate metric.

Mitotic cells displayed no changes in morphology prior to formation of daughter cells (*p* > 0.05 for all metrics). Daughter cells increased in size following division, while the circularity and instantaneous speed decreased. The nearest neighbors of mitotic cells displayed a decrease in circularity over the 60-minute period encompassing 30 minutes before and after cytokinesis. Apoptotic cells displayed loss of area and perimeter, as well as an increase in instantaneous speed. Interestingly, the nearest neighbors of apoptotic cells displayed the most substantial morphological changes highlighting their important role in maintenance of barrier. During apoptosis, the nearest neighbors displayed a reduction in area (*p* = 0.0144), showing that they did not contribute to compensation of the loss of monolayer area. This observation implies that more distant cells must increase in area and push the nearest neighbors into the area lost by the apoptotic cell. The extrusion of apoptotic cells has been reported for epithelial monolayers [54]. Additionally, the nearest neighbors of apoptotic cells displayed reduced circularity, number of cell neighbors (as a neighbor was lost by the end of apoptosis), and highly dynamic instantaneous speeds that were increased after the point of cell collapse but reduced at 30 minutes after apoptosis compared to 30 minutes before (*p* = 0.0117, < 0.0001, and 0.0249, respectively) (**Fig. S9**). The decrease in circularity of the nearest neighbors suggests that weakening the shared junction with the apoptotic cell leads to asymmetry and elongation perpendicular to the radial direction from the defect.

Having identified the characteristics of quiescent, mitotic, and apoptotic cells, and their nearest neighbors, we next compared morphological metrics between cell types before and after these events (**Fig. 4F-G, Fig. S10**). Prior to mitosis or apoptosis (*t* = - 30 minutes), cells generally displayed similar morphology, however, apoptotic cells and nearest neighbors to mitotic cells displayed lower instantaneous speed (*p* = 0.0459 and 0.0379, respectively), suggesting that changes in cell motility precede these events (**Fig. 4F**). Interestingly, neighbors of apoptotic cells displayed fewer neighboring cells themselves (*p* < 0.0001), suggesting that apoptosis occurred preferentially in local regions of low cell density. After apoptosis or mitosis (*t* = 30 minutes), daughter cells and nearest neighbors of apoptotic cells displayed distinct morphology compared to quiescent cells (**Fig. 4G, S8**). Daughter cells remained smaller and displayed lower instantaneous speed (*p* = 0.0003 and 0.0211, respectively), while nearest neighbors of apoptotic cells remained smaller and had fewer cell neighbors (*p* = 0.0064 and < 0.0001, respectively).

### iBMEC microvessel response to physical insult by laser ablation

After defining the hallmarks of iBMEC turnover during homeostasis, we next explored the response of microvessels to physical insult. Various processes can lead to vascular injury, including stroke (e.g. rupture/occlusion), neurodegenerative disease (e.g. cerebral amyloid angiopathy), or as a result of treatment (e.g. radiotherapy) [55]. Laser ablation was used to introduce small defects (5 - 10 cells) into iBMEC microvessels, resulting in Evans Blue extravasation at the wound site (**Fig. 5A**), similar to the extravasation of blood components observed *in vivo* following laser ablation of sub-cortical vessels [56]. The speed of wound healing was very similar between 2D and 3D (τ = ∼19 and ∼16 minutes, respectively). In 3D microvessels, completed defect recovery occurred over ∼2 h, during which we tracked wound area and the dynamics of individual cells bordering the wound (**Fig. 5B-C, Fig. S11A**). Following closure, cell debris was moved to the center of the defect forming a visible scar that persisted for a further 2 - 4 h (**Fig. 5B**), however, after 24 h there was no visible evidence of the wound. As there was no change in the rates of apoptosis, mitosis, and turnover between ablated microvessels and homeostatic controls (*p* = 0.414, 0.287, and 0.466, respectively), repair occurred through a combination of cell migration to the defect and an increase in area of the surrounding cells. Indeed, from cell tracking, we found that cells within ∼7 near neighbors of the wound displayed directed migration towards the wound centroid (**Fig. S11B-C**). This contrasts apoptosis where cells beyond the first nearest neighbors displayed negligible directed motion (**Fig. 5D**). Additionally, we observed an approximately 15% increase in average cell area after wound closure (p = 0.0010, Mann-Whitney test) (**Fig. 5E, Fig. S13A-B**). As wounds comprised on average ∼13% of the initial imaging frame, this confirms that compensation of lost area occurs predominately via increases in cell size, not proliferation. The number of the smallest cells per unit area (histogram bin: 50 – 150 µm^2^) was reduced by more than two-fold following wound closure, suggesting that primarily small cells increase their area to compensate for lost area from the monolayer (**Fig. 5E**).

**Figure 5.**
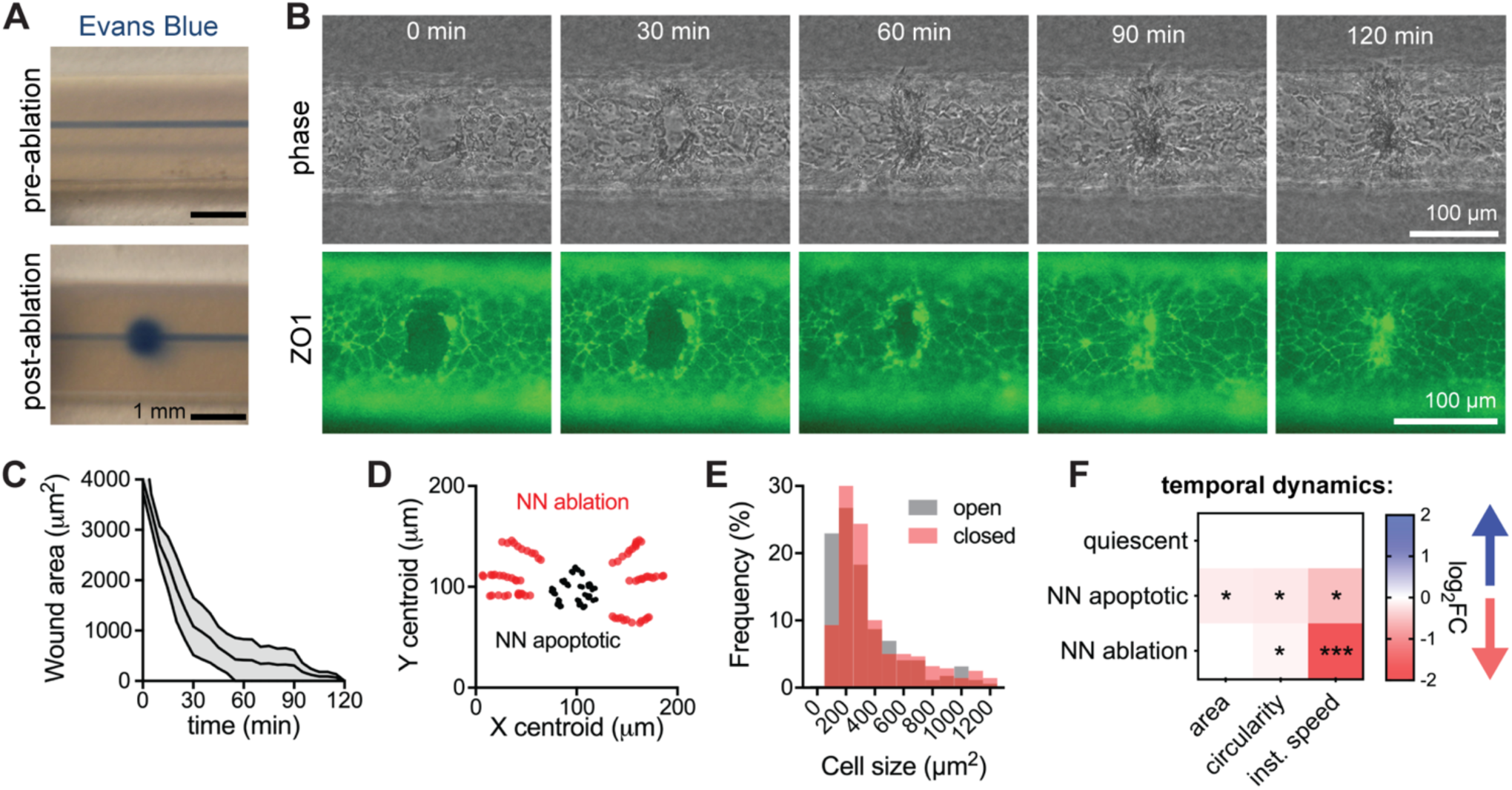
Response of three-dimensional iBMEC microvessels to physical injury. (A) Representative photographs of Evans Blue leakage induced by laser ablation of an area corresponding to 5 – 10 cells. (B) Live-cell imaging of TJ-labeled iBMEC microvessel recovery following laser ablation. (C) Time course of wound size (*n* = 4 microvessels). (D) Centroid tracking of cell neighbors surrounding a wound during healing (red dots), as well as cell neighbors of an apoptotic cell (black dots). Cell neighbors of ablation display directed migration towards the centroid of the wound. (E) Cell size distribution immediately after ablation and following wound closure of a total monolayer area of 35,000 µm^2^ (*n* = 353 cells in open wound and *n* = 290 cells in closed wound). (F) Heatmap comparing changes in cell morphology of nearest neighbors surrounding a wound during healing, relative to quiescent cells and neighbors of an apoptotic cell. Only squares corresponding to a statistically significance change are shaded by magnitude of change (log_2_FC) and direction of change (blue: increasing over time, red: decreasing over time) labeled with asterisks representing significance as calculated using the maximally significant result between a Wilcoxon matched-pairs signed rank test to compare first and last time points and using an F test to determine if linear regression of the morphological time course displayed a statistically non-zero slope. NN – nearest neighbors. Data are presented as mean ± SEM. * *p* < 0.05, ** *p* < 0.01, and *** *p* < 0.001. See also **Figure S11-13**; all analysis conducted on TJ-labeled iBMECs.

The cells immediately neighboring the wound displayed no change in area (p = 0.6095, Wilcoxon test), but did display decreasing speed (p < 0.0001, Wilcoxon test), indicating that repair is mediated at the monolayer level (**Fig. 5F, Fig. S12B-C**).

### iBMEC microvessel response to chemical injury

Diverse chemical perturbations are capable of modulating BBB function. To understand mechanisms of chemical injury, we exposed iBMEC microvessels to menadione and melittin. Menadione is a naturally occurring compound that can generate intracellular reactive oxygen species (ROS) and induce oxidative stress [57]; melittin is a membrane active peptide that can increase the permeability of cellular barriers [58]. While both menadione and melittin result in rapidly reduced TEER in Transwells, the mechanisms of barrier loss are not well characterized. Within 3D microvessels, we modeled chronic BBB injury by perfusion with 1 mM menadione and acute BBB injury by 10-minute exposure to 10 μM melittin (**Fig. 6A-B**). Using simultaneous phase contrast and fluorescence imaging we tracked microvessel structure, barrier properties, and tight junction dynamics.

**Figure 6.**
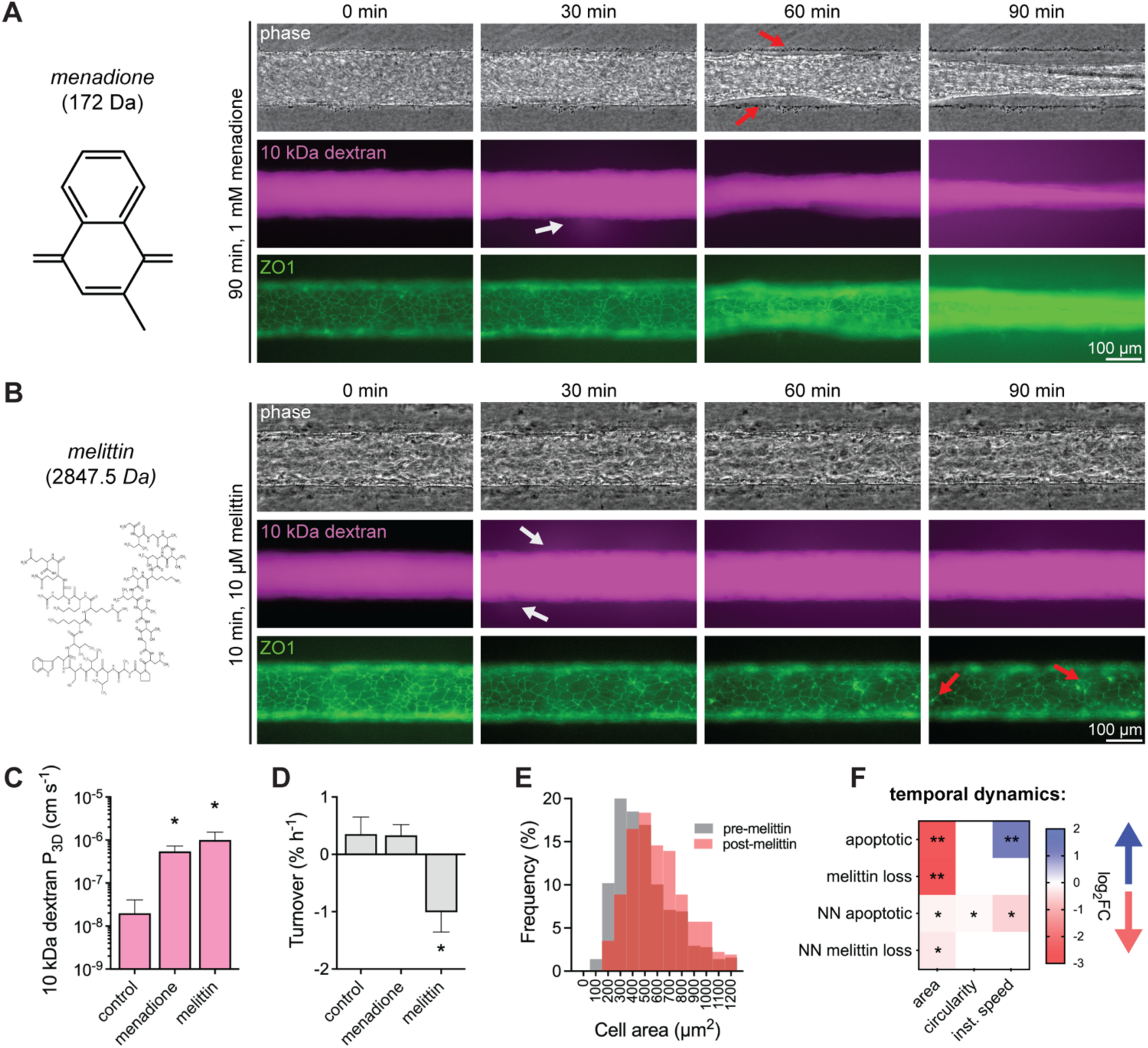
Response of three-dimensional iBMEC microvessels to chemical injury. (A-B) Real-time imaging of iBMEC microvessels in response to perfusion with menadione or melittin exposure. (A) Red arrows indicate delamination of the endothelium and white arrows denote sites of leakage of 10 kDa dextran. (B) Red arrows indicate cell collapse and white arrows denote sites of leakage of 10 kDa dextran. (C-D) Comparison of 10 kDa dextran permeability and cell turnover between control (*n* = 5), menadione-exposed (*n* = 4), and melittin-exposed (*n* = 4) iBMEC microvessels. Significance analysis performed using a Kruskal-Wallis test and post hoc Dunn’s multiple comparisons test relative to control. (E) Cell size distribution before and after melittin exposure (*n* = 581 cells pre-melittin and 329 cells post-melittin). (F) Heatmap comparing changes in cell morphology for apoptotic cells and cell loss induced by melittin exposure, as well as their respective nearest neighbors (NN). Only squares corresponding to a statistically significance change are shaded by magnitude of change (log_2_FC) and direction of change (blue: increasing over time, red: decreasing over time) labeled with asterisks representing significance as calculated using the maximally significant result between a Wilcoxon matched-pairs signed rank test to compare first and last time points and using an F test to determine if linear regression of the morphological time course displayed a statistically non-zero slope. NN – nearest neighbors. Data are presented as mean ± SEM. * *p* < 0.05, ** *p* < 0.01, and *** *p* < 0.001. See also **Figure S12-13**; all analyses conducted on TJ-labeled iBMECs.

Both menadione and melittin increased permeability of 10 kDa dextran (**Fig. 6A-C**) (*p* = 0.0318 and 0.0411), but with distinct modes of disruption: menadione induced delamination of the endothelium and partial collapse of the microvessel, whereas melittin induced cell loss from the endothelium (**Fig. 6A-B**). Endothelial turnover was dramatically increased by melittin (*p* = 0.0158), but not menadione exposure (*p* = 0.9584) (**Fig. 6D**). The increase in turnover in response to melittin was due to an ∼3-fold increase in cell loss (*p* = 0.0053) while the mitosis rate was unchanged (*p* = 0.3968). The cell loss due to melittin exposure resulted in a redistribution of cell area (**Fig. 6E, S13C-D**); before melittin exposure the cell area was 500 ± 10 μm^2^, while after melittin exposure the cell area increased to 641 ± 16 μm^2^ (*p* < 0.0001, Mann-Whitney test). In contrast to apoptotic cells, there was no change in instantaneous speed of lost cells in response to melittin, and no changes in circularity or instantaneous speed for neighbors of lost cells (**Fig. 6F**). Additionally, we tracked individual cell size and circularity finding that cells lost during melittin exposure had similar area (*p* = 0.7696), but lower circularity (*p* < 0.0001) compared to the entire monolayer (**Fig. S13E**). The average cell area in the monolayer was also observed to be predictive of melittin response (*r*^*2*^ = 0.9214) (**Fig. S13F**).

## Discussion

iPSC-derived BMEC-like cells (iBMECs) in 2D monolayers recapitulate aspects of BBB function, however, gene expression profiles suggest a component of epithelial identity. One approach to overcome this limitation is to reprogram iBMECs using ETS transcription factors. Although this results in gene expression profiles resembling primary, immortalized, and iPSC-derived endothelial cells (iECs), it results in significant loss of barrier function compared to iBMECs [23]. Thus, a grand challenge remains in achieving high transcriptomic similarity to human BMECs while maintaining key phenotypic characteristics of the BBB. Here, we explore how cues associated with the 3D microenvironment including direct cell-cell and cell-ECM interactions, shear stress, and cylindrical geometry augment iBMEC phenotype. We found that 3D microenvironment: (1) decreased paracellular permeability, (2) induced gene expression changes distinct from shear stress alone, including increased endothelial gene expression, and (3) enhanced angiogenic and cytokine responses. Additionally, using our 3D microvessel model we demonstrated physiological cell-cell interactions with pericytes and conducted real-time imaging of solute permeability and tight junction dynamics across a range of perturbations.

Within 3D microvessels, Lucifer yellow permeability was ∼ 2 × 10^−7^ cm s^-1^, matching permeability values in animal models [59], and approximately 10-fold lower compared to 2D models. Interestingly, improved barrier function in 3D was not associated with upregulation of tight junctions, suggesting that this effect is mediated by other changes in gene and protein expression or by experimental differences (see *Note S4*). Endothelial gene expression was enriched in 3D, while shear stress alone did not exert these effects [28]. An important functional implication of these changes is that 3D iBMEC microvessels displayed bFGF-induced angiogenic sprouting and TNFα-induced immune cell adhesion, which have been inconsistently reported in 2D iBMEC models. GSEA suggested other unique phenotypes of iBMECs in 3D, which were distinct from the effects of shear stress. Although iBMECs in 3D microvessels exhibit some functional similarities to the human BBB, they still possess a component of epithelial identity (*EPCAM, CLDN6, CLDN3, CDH4*) and express lower levels of several endothelial transcripts (*PECAM1, CLDN5, TIE1, CDH5*) than primary, immortalized, and iPSC-derived endothelial cells (**Fig. S6**). However, the increased barrier strength of iBMECs (compared to primary and immortalized BMECs) cannot only be ascribed to epithelial identity, as MDCK epithelial cells, which have been the workhorse in BBB research since the 1970s, exhibit low TEER values (typically around 200 Ω cm^2^) [60]. 3D microenvironment was not found to change epithelial gene expression, while overexpression of ETS TFs to promote endothelial identity does dramatically downregulate epithelial transcripts [23]. Thus, synergistic approaches using co-cultured cell types, microenvironmental cues, and brain-specific TF overexpression will likely be key to improved physiological relevance of BBB models.

In addition to providing cues for promoting BBB phenotype, incorporating iBMECs into a 3D microenvironment enables real-time imaging of a diverse range of processes at the single-cell level. From live-cell imaging of tight junctions, we were able to study the cooperative behavior of iBMECs in confluent monolayers. While traditional techniques to probe tight junction dynamics (e.g. immunocytochemistry) display poor temporal resolution, we were able to map tight junction dynamics during all stages of turnover and in response to physical and chemical insult. Additionally, simultaneous measurements of permeability with high spatial resolution enabled direct correlation to changes in tight junction dynamics. The processes of apoptosis and mitosis were associated with distinct morphological changes of the cell as well as its neighboring cells, where the underpinnings of these cellular changes were distinct during physical and chemical injury. Notably, the turnover dynamics (**Fig. S12A**) and response of neighbors of cells lost from the endothelium (**Fig. S12B-C**) were unique across modes of injury. Lost area due to apoptosis in homeostatic microvessels was not directly compensated by increased area of neighboring cells, while during wound healing loss of area was compensated by an increase in area of surrounding cells with small initial area. Contrastingly, in response to melittin exposure there was a dramatic increase in cell area across the monolayer which was associated with cell loss and increased paracellular leakage.

3D models also enable incorporation of other cell types with physiological cell-cell interactions that are not possible in Transwell systems. Previously, we have shown that co-culture of pericytes in 3D iBMEC microvessels does not change solute permeability [61]. However, here we observed that pericyte co-culture modulated other phenotypes including angiogenic and cytokine responses. Matching *in vivo* observations [62], we found that pericytes increased the density of angiogenic sprouts from Ibmec microvessels. While *in vivo* work in pericyte-deficient mice finds that pericytes limit leukocyte infiltration in the absence of TNFα exposure [47], here, we found that pericytes increase the magnitude of TNFα-induced immune cell adhesion, while maintaining low levels of adhesion under baseline conditions. These results suggest that pericytes can alter properties of the BBB beyond barrier function, and may guide when pericyte co-culture is critical for *in vitro* applications.

In summary, through comparison of gene expression and functional properties of iBMECs in 2D monolayers and in 3D tissue-engineered microvessels, we show how microenvironmental cues promotes BBB phenotype, including enhanced endothelial identity, angiogenic response, and cytokine response, which extends the repertoire of experiments possible using iBMECs. Despite the mixed endothelial and epithelial identity, iBMECs in 3D microenvironments provide a powerful tool for studies of the BBB, and could in the future be combined with other chemical or transcription factor overexpression approaches. Specifically, we applied iBMECs in 3D microvessels to enable detailed visualization and analysis of tight junction dynamics during homeostasis, wound repair, and chemical injury. Our results present new insight into how monolayers of brain microvascular endothelial cells cooperatively respond to mitosis and apoptosis events, as well as to physical or chemical insult.

## Methods

### Cell culture and characterization

BMEC-like cells (iBMECs) were differentiated based on published protocols (see *Supplemental Experimental Procedures*) [22, 52] from three isogenic WTC iPSC lines (Allen Cell Institute) [63]: enhanced green fluorescence protein (EGFP)-labeled zona occludens-1 (ZO1) (iPSC-TJ), red fluorescence protein (RFP)-labeled plasma membrane (iPSC-PM), and EGFP-labeled β-actin (iPSC-ACTB). These iPSCs correspond to cell line IDs: AICS-0023 cl.20, AICS-0054 cl.91, and AICS-0016 cl.184, respectively. Additional experiments were conducted on non-isogenic BC1 iBMECs [22, 52]. Differentiations in which peak TEER values were below 1500 Ω cm^2^ (< 10% frequency) were excluded from analysis. Isogenic iPSC-derived endothelial cells (iECs) were differentiated using *ETV2* modRNA following published protocols [40]. The iECs were cultured in endothelial cell growth medium 2 kit supplemented into basal media (except hydrocortisone, Lonza) with 1x GlutaMax (ThermoFisher) and 10 μM SB431542 (Selleckchem). Brain pericyte-like cells (iPCs) were derived through a neural crest intermediate using published protocols (see *Supplemental Methods*) [49]. The iPCs were cultured in E6 media (StemCell Technologies) supplemented with 10% FBS and routinely passed using Accutase at 1:5 on tissue-cultured treated surfaces. Primary neonatal human dermal microvascular endothelial cells (HDMECs: Lonza) were cultured in MCDB 131 (Caisson Labs, Carlsbad, CA) supplemented with 10% heat inactivated fetal bovine serum (Sigma), 25 mg mL^-1^ endothelial mitogen (Biomedical Technologies), 2 U mL^-1^ heparin (Sigma), 1 μg mL^-1^ hydrocortisone (Sigma), 0.2 mM ascorbic acid 2-phosphate (49752, Sigma), and 1% penicillin-streptomycin-glutamine (ThermoFisher). The details of cell culture, differentiation, barrier assays, and recordings of glucose and oxygen levels during iBMEC differentiation are provided in *Supplemental Methods*.

### Microvessel fabrication and imaging

3D microvessels were fabricated similar to previously reported methods [52]. Briefly, 150 μm diameter channels were patterned in (1) 7 mg mL^-1^ collagen I crosslinked with 20 mM genipin, or (2) 6 mg mL^-1^ collagen I and 1.5 mg mL^-1^ Matrigel. Channels were connected to inlet and outlet ports within a PDMS-based microfluidic device to control flow rates and shear stress, with perfusion maintained by a gravity driven flow system achieving average flow rates of ∼0.25 mL h^-1^. iBMECs (differentiated using 1 mL media protocols) and iECs were seeded into the channels and allowed to adhere for 30 minutes before initiating perfusion. To form co-cultured microvessels, iPCs were seeded into the channels and perfused at 1 dyne cm^-2^ for 24 hours prior to seeding iBMECs. 40x confocal images were obtained using a swept field confocal microscope system (Prairie Technologies) with illumination provided by an MLC 400 monolithic laser combiner (Keysight Technologies). 10x epifluorescence images were obtained using an inverted microscope (Nikon Eclipse Ti-E), with illumination was provided by an X-Cite 120LEDBoost (Excelitas Technologies). Time lapse images were acquired every 2 or 5 minutes depending on the experiment, in an environmental chamber maintained at 37 °C and 5% CO_2_.

Fluorescence images at the microvessel midplane were acquired every two minutes before (10 minutes total) and after solute perfusion (60 minutes total). Filter cubes (Chroma 39008 and Chroma 41008) were used to independently capture Lucifer yellow (20 ms exposure) and Alexa Fluor-647-conjugated dextran (200 ms exposure), used at concentrations matching 2D assays (see *Supplemental Methods*). Images were collected as ten adjacent frames corresponding to a total image area of 8.18 mm x 0.67 mm. ImageJ was used to plot fluorescence intensity profiles over 70 minutes (36 frames). Permeability (P) = (r/2)(1/ΔI)(dI/dt)_0_, where r is the microvessel radius, ΔI is the increase in fluorescence intensity due to luminal filling, and (dI/dt)_0_ is the rate of fluorescence intensity increase (calculated over 60 minutes) [64].

Images were segmented into ten adjacent regions-of-interest (ROIs), where the permeability is reported as the mean value of the five adjacent frames surrounding the minimum to minimize artifacts from interstitial dye entering the matrix from inlet and outlet ports during imaging. For details of permeability calculations see *Note S4*.

### Immunocytochemistry

For 2D experiments, iBMECs were seeded at 250,000 cells cm^-2^ on borosilicate cover glass slides (Thermo Scientific); for 3D experiments, iBMECs were seeded as described above. Two days later, cells were washed with phosphate-buffered saline (PBS; ThermoFisher), fixed with ice-cold methanol for 15 minutes, and blocked with 10% goat serum (Cell Signaling Technology) or 10% donkey serum (Millipore Sigma), supplemented with 0.1% Triton X-100 (Millipore Sigma) for 30 minutes. Primary antibodies were exposed overnight at 4C with details listed in **Table S2**. After three washes with PBS, cells were treated with Alexa Flour-647 and Alexa Flour-488 secondary antibodies (Life Technologies) diluted 1:200 in blocking buffer for 1 hour at room temperature. Nuclei were stained using 1 μg mL^-1^ DAPI (ThermoFisher). Confocal images were acquired at 40x magnification as previously reported [52]. Control images were also collected without primary antibody to confirm fluorescence above non-specific background. Semi-quantitative analysis of protein levels between 2D and 3D was calculated based on the ratio of immunofluorescence between 3D and 2D, and normalized to the ratio of nucleus fluorescence (by DAPI staining) between 3D and 2D, and normalized by endothelial surface area.

### Bulk RNA sequencing

RNA was collected two days after subculture or passaging. Cells were washed with PBS and lysed using RLT buffer supplemented with 1% β-mercaptoethanol. Lysates were eluted with RNase-free water after purification using a RNeasy Mini Kit (Qiagen) and DNase I digestion, following manufacturer instructions. All samples had an RNA integrity number > 8.4 as measured by an Agilent 2100 bioanalyzer. Total RNA was subjected to oligo (dT) capture and enrichment, and the resulting mRNA fraction was used to construct cDNA libraries (performed by Novogene). Sequencing was carried out on an Illumina NovaSeq platform (performed by Novogene) with paired end 150 bp reads, generating approximately 20 million paired reads per sample. The R (v4.0.1) package Rsubread (v2.0.1) was used for raw read alignment and for read quantification to the reference human genome (GRCh38) [77]. The R package DESeq2 (v1.28.1) was used for normalization, visualization, and differential analysis [78]. Raw reads were normalized using the DESeq2 variance stabilizing regularized logarithm (rlog) transformation prior to calculation of Euclidean sample distances, Spearman correlation coefficients, and principal component analysis (PCA). Differentially expressed genes (DEGs) were determined using the Wald test with Benjamini-Hochberg correction, where adjusted *p* values < 0.05 was considered statistically significant. Pathway enrichment analysis was conducted via genome-wide ranked list comparisons using Gene Set Enrichment Analysis (GSEA, v4.1.0) for the Molecular Signatures Database (MSigDB) hallmark gene sets with 1000 permutations and a false discovery rate < 0.25; normalized enrichment score (NES) was calculated by the software [44, 79]. Plots were all formatted using the R packages ggplot2 (v3.3.2), ggrepel (v0.8.2), pheatmap (v1.0.12), and RColorBrewer (v1.1-2). Published bulk transcriptomes were obtained from the NCBI Gene Expression Omnibus (GEO) (see Table S1).

### Quantifying angiogenic and cytokine response

To probe angiogenic response, microvessels with and without pericytes were formed on 6 mg mL^-1^ collagen I and 1.5 mg mL^-1^ Matrigel hydrogels. As a static control, microspheres were seeded with iBMECs and embedded within hydrogels as previously reported [50]. After 48 h treatment with 20 ng mL^-1^ bFGF (R&D Systems), sprouts were manually counted in ImageJ and sprout length calculated as the total length of sprouts per unit area of endothelium.

To probe cytokine responses, microvessels with and without pericytes were formed in 7 mg mL^-1^ collagen I crosslinked with 20 mM genipin. THP-1 (ATCC® TIB-202™), a human leukemia monocytic cell line [65], were grown in suspension with RPMI-1640 Medium (Sigma) supplemented with 10% fetal bovine serum (Sigma) and 1% penicillin-streptomycin. Cells were labeled with 1 μM CellTracker™ Red CMTPX Dye (ThermoFisher) in serum-free media for 20 min, and then resuspended at 1 × 10^6^ cells mL^-1^ in complete media. Microvessels with and without 24 h treatment with 5 ng mL^-1^ human recombinant TNFα (R&D Systems), were perfused with THP1s under low shear stress (∼0.2 dyne cm^-2^) for 10 min, and then washed out using higher shear stress (∼2 dyne cm^-2^). As a 2D control, iBMECs monolayers were exposed to THP1s matching microvessel conditions, with washout mimicked by conducting three gentle media washes. Adherent immune cells were manually counted in ImageJ and normalized to unit area of endothelium. Bulk RNA was extracted from microvessels 24 h after 5 ng mL^-1^ TNFα exposure.

### Quantifying tight junction dynamics

Images were collected every 5 minutes of Hoescht-33342 (ThermoFisher)-labeled nuclei, EGFP-labeled ZO1, and phase contrast. Endothelial cell dynamics within the monolayer were quantified using previously developed tools in ImageJ (NIH) [52, 66]. From a total of 1,871 cells, all apoptotic and mitotic cells with 30 minutes of imaging before and after these events were included in analysis; control cells were randomly selected. Rates of apoptosis, mitosis, and turnover are not statistically significantly different between measurements based on phase contrast (as previously utilized [27]) or ZO1 fluorescence (*p* > 0.05 for all comparisons). Using ZO1 traces, cell area (µm^2^), perimeter (µm), circularity, aspect ratio, and centroid location of individual cells were then calculated from freehand tracings of cell boundaries. The number of nearest cell neighbors were manually counted based on based edges of tight junction expression. Instantaneous cell speed (nm s^-1^) was calculated from the change in the location of the centroid of the cell divided by elapsed time. The cell speed in the x-direction (along the length of the vessel and in the direction of flow) was determined from the total change in position in the x-direction divided by elapsed time. Results were validated across independent researchers to confirm robustness (**Fig. S6D**). PCA was conducted using the “prcomp” function in the R stats package (v 3.6.2). Metrics were first shifted to be zero-centered and scaled to have unit variance. Contributions of morphological metrics to principal components 1 – 4 were quantified using the squared cosine metric (**Fig. S6C**).

### Modeling physical and chemical injury

Laser ablation was conducted via 90 s irradiation at 2100 mW using the 750 nm line of an LSM 510 laser scanning microscope (Zeiss), where a rectangular prism (50 µm width x 150 µm length x 20 µm height) at the bottom plane of microvessels was irradiated. For 2D studies, iBMEC monolayers were scratched with a pipette tip to achieve a similar defect area. The defect area (µm^2^) was determined by tracing the perimeter of the wound at each time point. Barrier function was also assessed following perfusion with 0.0683% w/v Evans blue (in medium), matching concentrations in the bloodstream of adult mice dosed at 1% weight/volume [67]. To model chemical injury, microvessels were perfused with 1 mM menadione (Sigma) for 90 minutes or 10 μM melittin with a free carboxy C-terminus (melittin-COO^-^) (Bio-Synthesis Inc) for 10 minutes. Simultaneously, microvessels were perfused with 10 kDa dextran with phase and fluorescence imaging conducted over 90 minutes.

### Statistical analysis

Statistical testing was performed using Prism ver. 8 (GraphPad). All experimental values are reported as mean ± standard error of the mean (SEM). Statistical tests were chosen based on paired status and normality (Shapiro-Wilk test), where details are provided in figure captions. Differences were considered statistically significant with the following thresholds: * p < 0.05, ** p < 0.01, *** p < 0.001, and **** p < 0.0001.

## Data and materials availability

All data associated with this study are available in the main text or the supplementary materials. RNA sequencing data are deposited in GEO under accession number GSE195519.

## Acknowledgements

The authors also acknowledge the assistance of Alanna Farrell and Erin Pryce. This work was supported by DTRA (HDTRA1-15-1-0046) and NIH (R01NS106008 and R61HL154252). RML acknowledges a National Science Foundation Graduate Research Fellowship under Grant No. DGE1746891 and the support of Jeffrey Herman during long breaks from the benchtop during the COVID-19 pandemic.

