## Supplemental Information for "Three-dimensional microenvironment regulates gene expression, function, and tight junction dynamics of iPSC-derived blood-brain barrier microvessels"

**Supplemental Figures:**
Figure S1. Characterization of iBMEC phenotype following differentiation in 1 mL or 2 mL of medium.
Figure S2. TEER and permeability measurements of 2D iBMEC confluent monolayers.
Figure S3. Media volume effect on phenotype of 2D iBMEC confluent monolayers.
Figure S4. Details of bulk RNA sequencing and iPC characterization.
Figure S5. Benchmarking iBMEC gene expression to brain microvessel, endothelial, and epithelial datasets.
Figure S6. Abundance measurements of endothelial and epithelial transcripts across cell sources and model types.
Figure S7. Images of angiogenic sprouts across models.
Figure S8. Assessment of tight junction dynamics in confluent iBMEC monolayers.
Figure S9. Time course of morphological metrics across cell types.
Figure S10. Violin plots of metrics across cell types.
Figure S11. Response of three-dimensional iBMEC microvessels to wound formation by laser ablation.
Figure S12. Comparison of morphological metrics between homeostasis, ablation, and melittin exposure.
Figure S13. Monolayer area dynamics during wound healing and melittin exposure.

**Supplemental Tables:**
Table S1. Summary of bulk RNA transcriptomes used in this study.
Table S2. Antibodies used in this study.

**Supplemental Data Sets:**
Data S1. Summary of bulk RNA-sequencing data.
Data S2. Summary of tight junction dynamics data.

**Supplemental Notes:**
Note S1. Validation of media volume effect on TEER
Note S2. Variability of iBMEC differentiation
Note S3. Mechanisms of media volume effect on iBMEC phenotype
Note S4. Calculation of permeability in 2D and 3D.

**Supplemental Methods:**
*iBMEC differentiation*
*Transwell barrier characterization*
*Oxygen and glucose recordings*
*Pericyte differentiation and characterization*

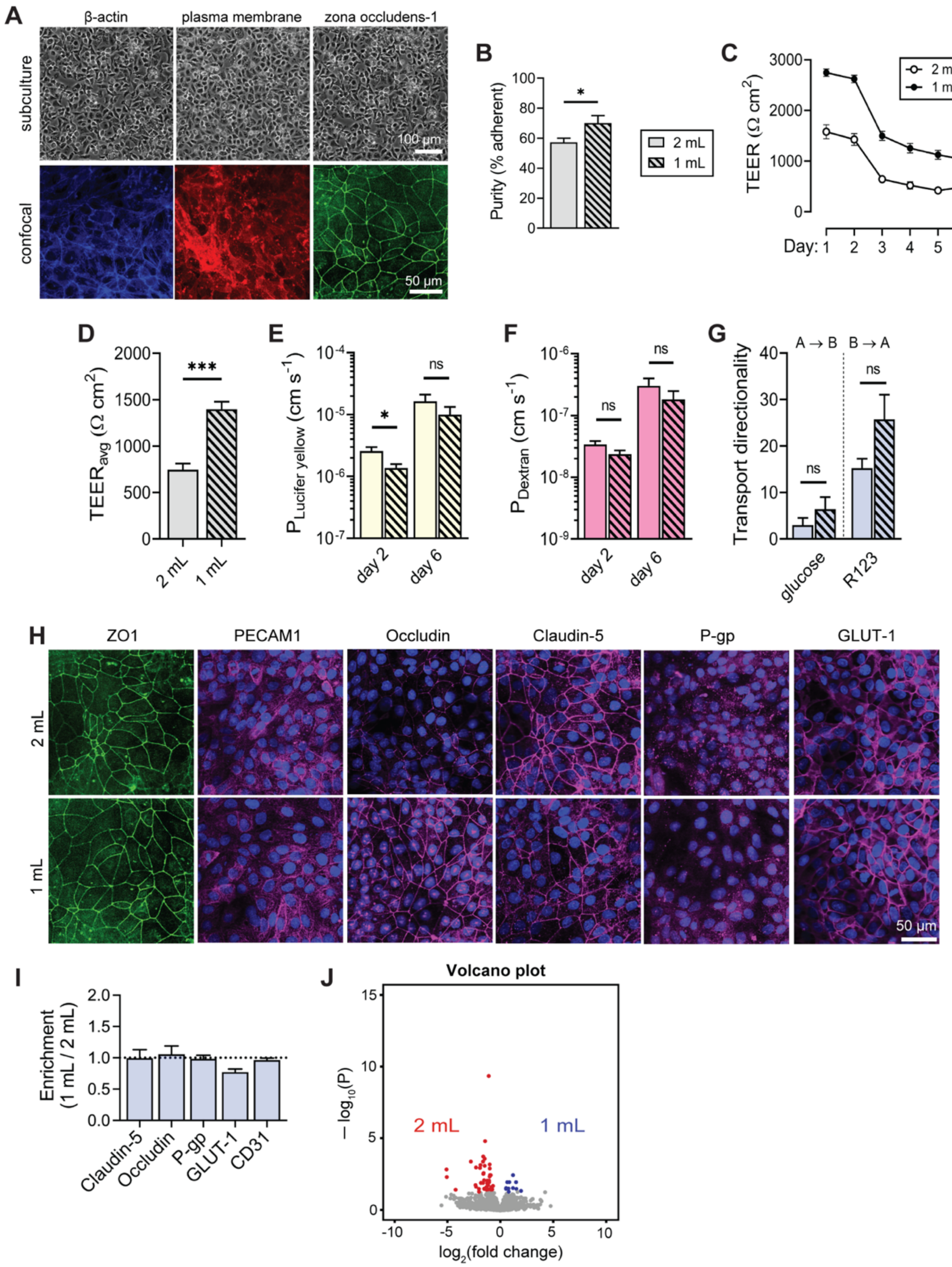


**Figure S1.** Characterization of iBMEC phenotype following differentiation in 1 mL or 2 mL of medium.

(A) Representative phase contrast images after sub-culture and confocal fluorescence images two days after subculture of iBMEC-TJs (tight junction, ZO-1), iBMEC-PMs (plasma membrane), and iBMEC-ACTBs (actin cytoskeleton).

(B) Purity of adherent cells emerging from differentiation using 2 mL or 1 mL media volumes ($n$ = 10 unique differentiations)

(C-D) Time course and average TEER between 1 mL and 2 mL differentiation ($n$ = 55 unique differentiations). Fluorescent-labeling does not alter barrier properties as similar TEER values were observed across all three cell lines.

(E-F) Comparison of Lucifer yellow and 10 kDa dextran permeability between media volumes ($n$ = 11 unique differentiations, measured on days 2 and 6).

(G) Glucose and rhodamine 123 transport is similar across differentiation media volumes ($n$ = 4 and 10 unique differentiations, respectively). Glucose is preferentially transported in the apical-to-basolateral direction (A → B), and hence directionality is defined as $P_{A \rightarrow B} / P_{B \rightarrow A}$. R123 is preferentially transported in the basolateral-to-apical direction (B → A), and hence directionality is defined as $P_{B \rightarrow A} / P_{A \rightarrow B}$.

(H) Localization of endothelial markers (CD31, VE-cadherin), tight junction proteins (ZO1, occludin, claudin-5), nutrient transporters (GLUT1), and efflux transporters (P-gp). Localization and expression levels are similar for 1 mL and 2 mL of media. Representative images are shown.

(I) Semi-quantitative analysis of protein expression between iBMECs cultured using 1 mL or 2 mL of media ($n$ = 3 – 9 unique differentiations per marker with values representing normalized fluorescence intensity).

(J) Volcano plots showing significantly (adjusted $p < 0.05$, Wald test with Benjamini-Hochberg correction) upregulated genes (blue) and downregulated genes (red) between 2D confluent monolayers iBMECs differentiated in 1 mL or 2 mL of media ($n$ = 3).

Data are presented as mean ± SEM. Statistical analysis was performed using a two-tailed paired t-test for two groups or a one-way ANOVA with Tukey multiple comparisons test for four groups: * $p < 0.05$, ** $p < 0.01$, and *** $p < 0.001$. See **Note S1**.

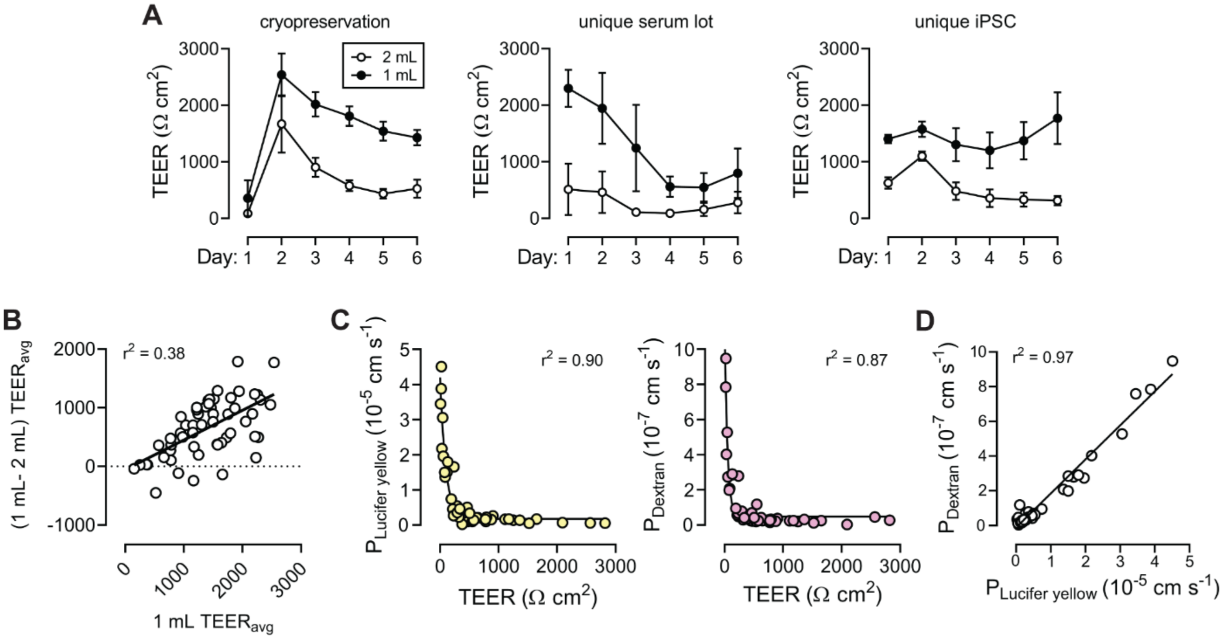


**Figure S2**. TEER and permeability measurements of 2D iBMEC confluent monolayers.

(A) TEER values for iBMEC monolayers are higher following differentiation in 1 mL medium. (1) Following cryopreservation (*n* = 4 paired independent differentiations), (2) using an independent serum lot (human serum from platelet poor human plasma lot #SLBT4321 and #SLCC2311 were used, where lot #SLCC2311 results are shown above) (*n* = 3 unique differentiations), and (3) using an independent iPSC source (BC1s) (*n* = 5 unique independent differentiations).

(B) Bland Altman plot of average TEER difference between differentiation media volumes versus average TEER for iBMECs differentiated using 1 mL (*n* = 55 unique differentiations). The solid line shows a linear least squares fit.

(C-D) Relationship between permeability and TEER. Permeability versus TEER fits to a one-phase exponential decay for both solutes ($r^2 > 0.86$). Lucifer yellow and 10 kDa dextran permeability are correlated linearly ($r^2 = 0.97$). Data collected across *n* = 48 independent Transwell measurements.

Data are presented as mean ± SEM.

*Summary*. Higher TEER values are obtained following differentiation in 1 mL medium, independent of cryopreservation, serum lot, or iPSC source. The permeability of Lucifer yellow and 10 kDa remains low for TEER values above 500 and 100 Ω $cm^2$, respectively. See **Note S2**.

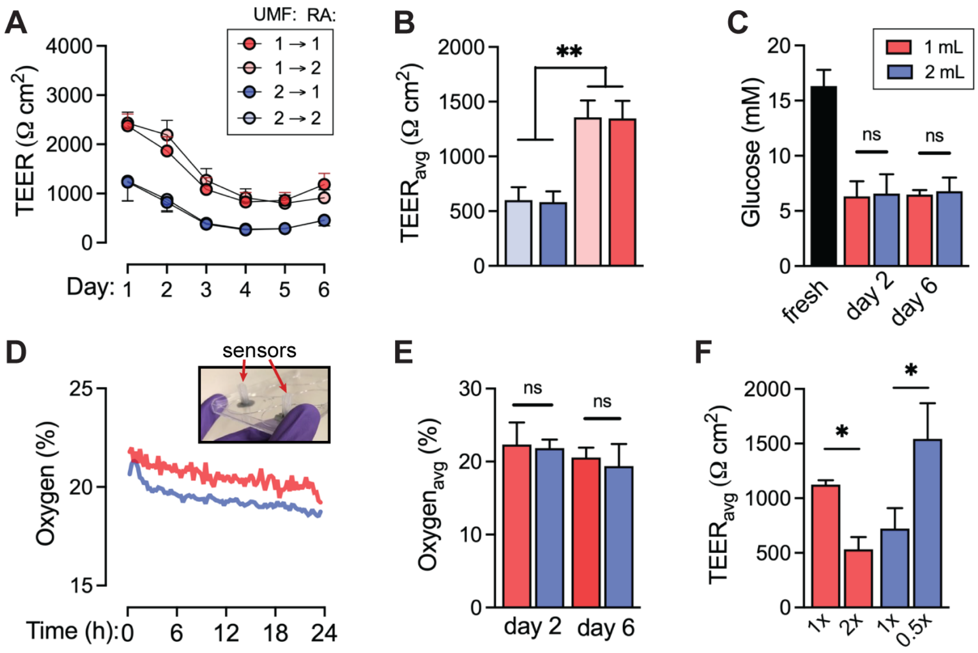


**Figure S3.** Media volume effect on phenotype of 2D iBMEC confluent monolayers.

(A-B) TEER time course and average TEER over one week for four conditions, using 1 mL or 2 mL at the UMF- and RA media phases ($n$ = 10 unique differentiations).

(C) Media glucose levels in fresh media compared to 1 mL and 2 mL medium at days 2 and 6, control ($n$ = 3 unique differentiations).

(D-E) Time course and average recordings of oxygenation between media volumes at day 2 and 6 of the differentiation. Inset shows sensors held in place on the bottom surface of the tissue culture plate ($n$ = 3 unique differentiations).

(F) Average TEER over one week for differentiations with altered media switch frequency (1 mL: $n$ = 3 unique differentiations, 2 mL: $n$ = 5 unique differentiations). In the standard differentiation, the UMF- medium is switched daily (6 days). There were no medium switches during the RA phase (2 days). 2x denotes switches of the UMF- twice daily, and 0.5x denotes UMF- medium switches every day.

Data are presented as mean ± SEM. Statistical analysis was performed using a two-tailed paired t-test for two groups or a one-way ANOVA with Tukey multiple comparisons test for four groups: * $p < 0.05$, and ** $p < 0.01$.

*Summary*. The highest TEER values were obtained when cells were differentiated in 1 mL medium in the UMF- phase, independent of the volume in the subsequent RA phase. Results show no significant difference in glucose or oxygen levels in 1 mL or 2 mL medium during differentiation. In general, higher TEER values were obtained with reduced medium switches during UMF and RA phases suggesting that barrier function is enhanced by secreted soluble factors or differences in cell-cell interactions during differentiation. See **Note S3**.

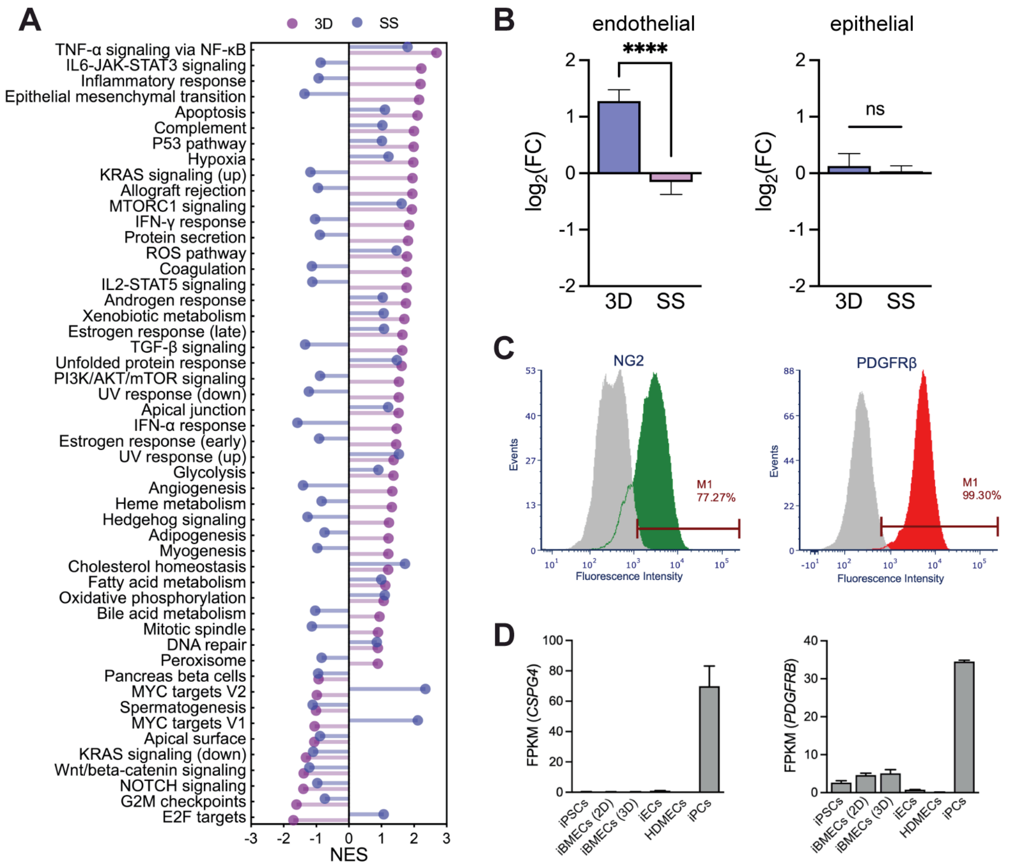


**Figure S4.** Details of bulk RNA sequencing and iPC characterization.

(A) Complete gene set enrichment analysis (GSEA) of hallmark gene sets.

(B) Distribution of $\log_2$FC for endothelial and epithelial transcripts between: (1) 3D versus 2D, and (2) flow versus static.

(C, D) Flow cytometry and RNA sequencing validation of iPCs. Expression of PDGFRβ and NG2 were confirmed at the protein and gene level.

Data are presented as mean ± SEM. **** $p < 0.0001$.

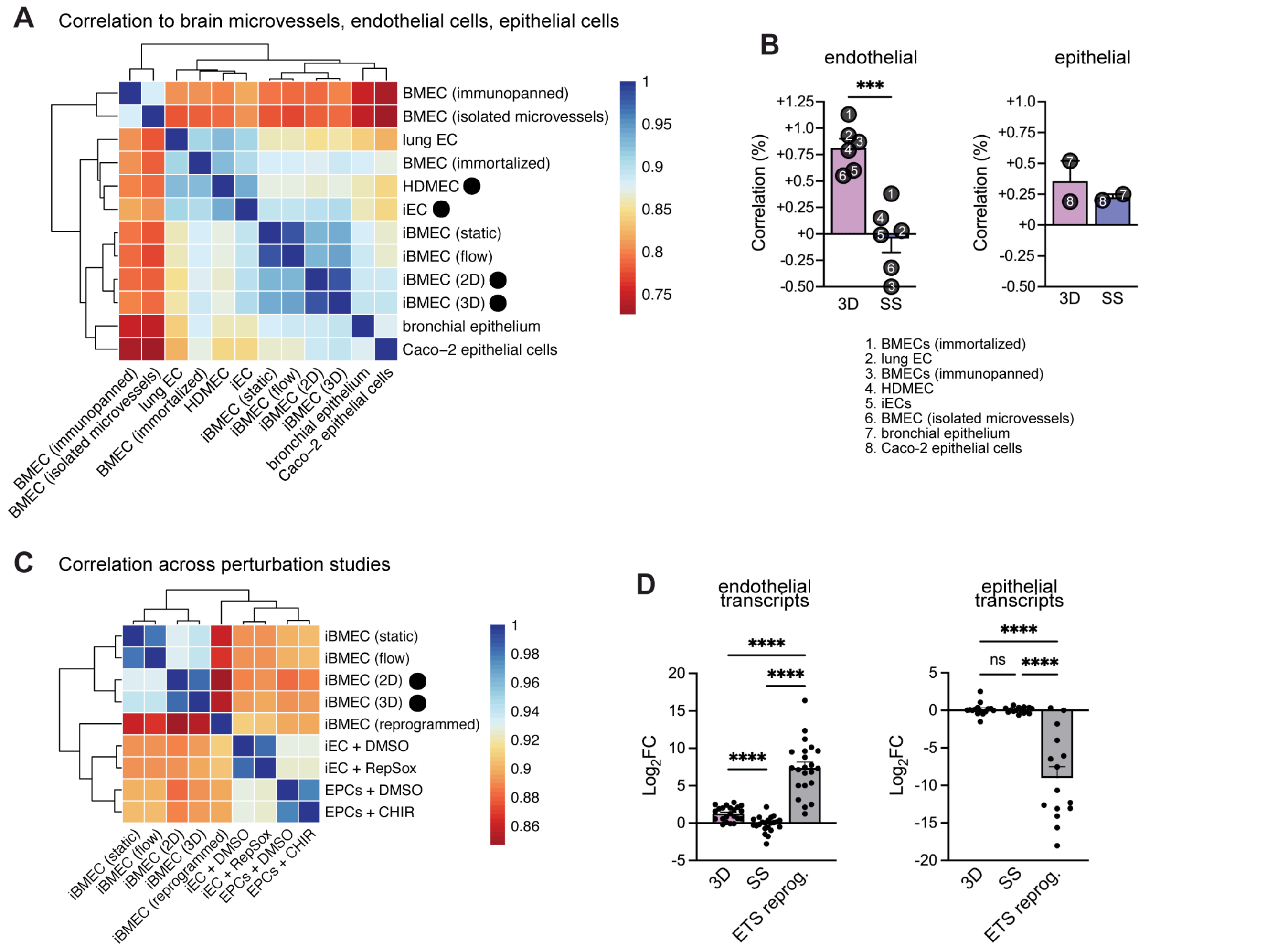


**Figure S5.** Benchmarking iBMEC gene expression to brain microvessel, endothelial, and epithelial datasets. iBMEC 2D and 3D (this work). See Table S1 for details of other data sets.

(A) Heatmap of spearman correlation coefficients across bulk RNA transcriptomes of brain microvessel, endothelial cells, and epithelial cells. Values determined from variance stabilized abundance measurements averages across samples belonging to the same experimental condition.

(B) Changes in iBMEC correlation to endothelial and epithelial datasets due to 3D microenvironment and shear stress.

(C) Heatmap of spearman correlation coefficients across bulk RNA transcriptomes of *in vitro* studies exploring methods to drive brain endothelial identity.

(D) Distribution of $\log_2$FC for endothelial and epithelial transcripts between: (1) 3D versus 2D, and (2) flow versus static, and (3) ETS TF reprogramming of iBMECs.

Data are presented as mean ± SEM. Black circles in S5A,C represent samples sequenced for this study. *** $p < 0.001$, **** $p < 0.0001$.

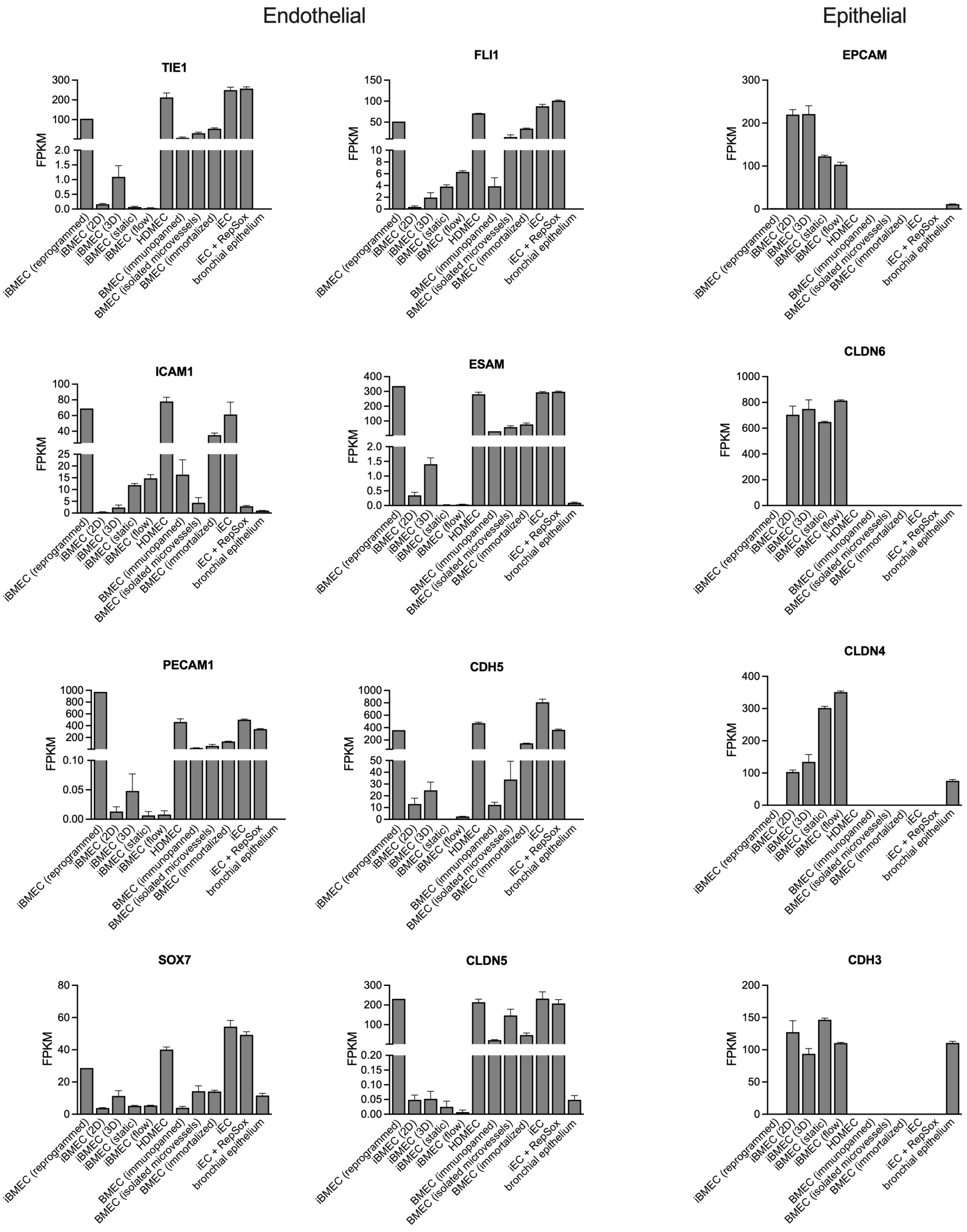


**Figure S6.** Abundance measurements of endothelial and epithelial transcripts across cell sources and model types.

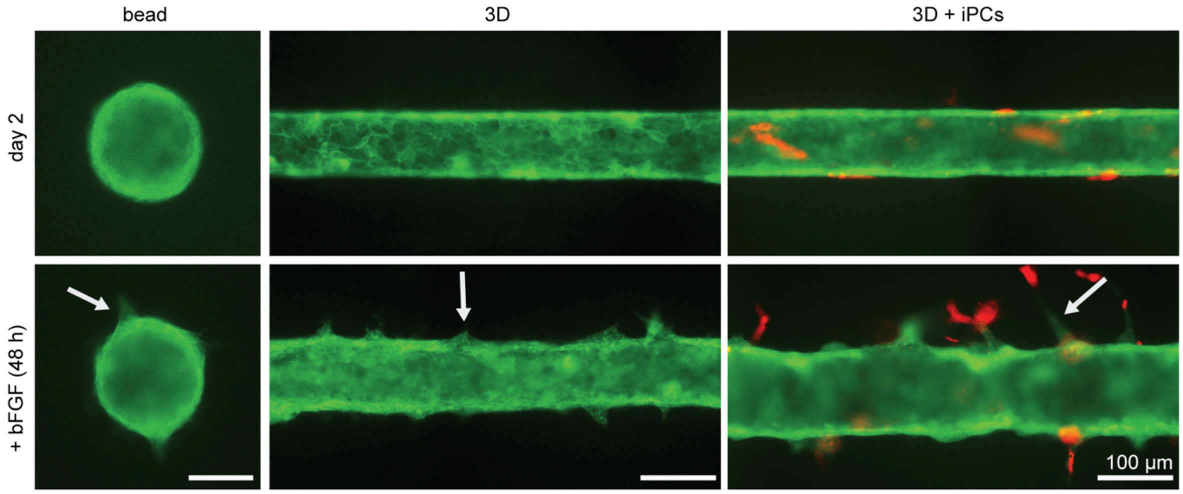


**Figure S7**. Representative images of iBMEC response to bFGF in iBMEC and iBMEC/iPC microvessels. Angiogenic sprouts appeared over two days of exposure to growth factor (white arrows). The isogenic co-culture BBB model was formed with ACTB-labeled iBMECs and PM-labeled iPCs. The microvessels were formed via sequential seeding of PM-labeled iPCs and then ACTB-labeled iBMECs, to achieve a 1:3 ratio (pericyte to EC). Pericytes were localized to the abluminal surface of the microvessel. Following maturation, microvessels were perfused for 48 h with 20 ng $mL^{-1}$ bFGF resulting in sprout formation.

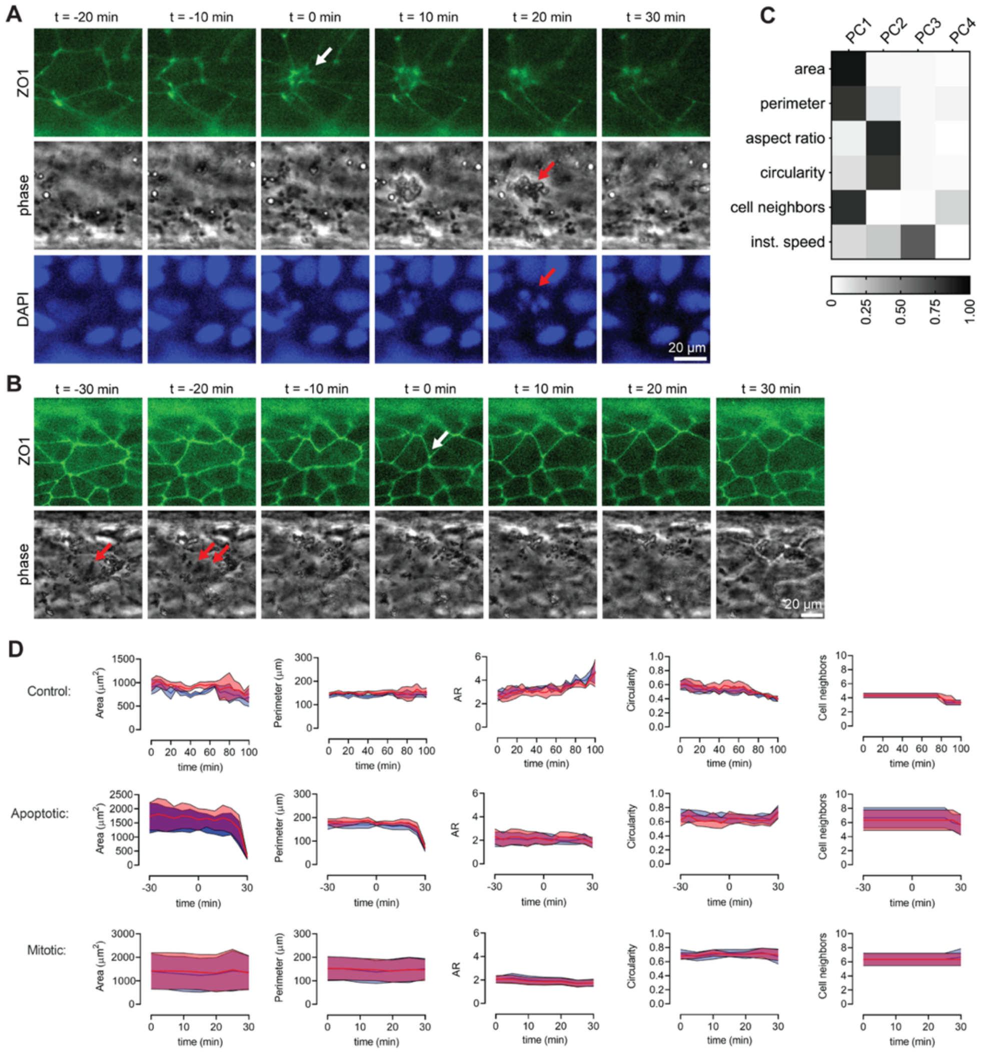


**Figure S8**. Assessment of tight junction dynamics in confluent iBMEC monolayers.

(A) Cells categorized as apoptotic display characteristic features in phase contrast images and nucleus fluorescence staining. Following cell collapse (white arrow), nuclear fragmentation and formation of apoptotic bodies are observed (red arrows).

(B) Cells categorized as mitotic display chromosomal alignment 30 minutes prior to formation of daughter cells (metaphase; red arrow), then separation of chromosomes (anaphase; two red arrows), then formation of new tight junctions (cytokinesis; white arrow).

(C) Principal component analysis highlights primary contributions to cell morphology (squared cosine).

(D) To validate assessment of morphological metrics, images were analyzed by two independent reviewers. Three cells for each classification were traced by reviewer 1 (red) and reviewer 2 (blue). Morphological metrics from both reviewers were similar.

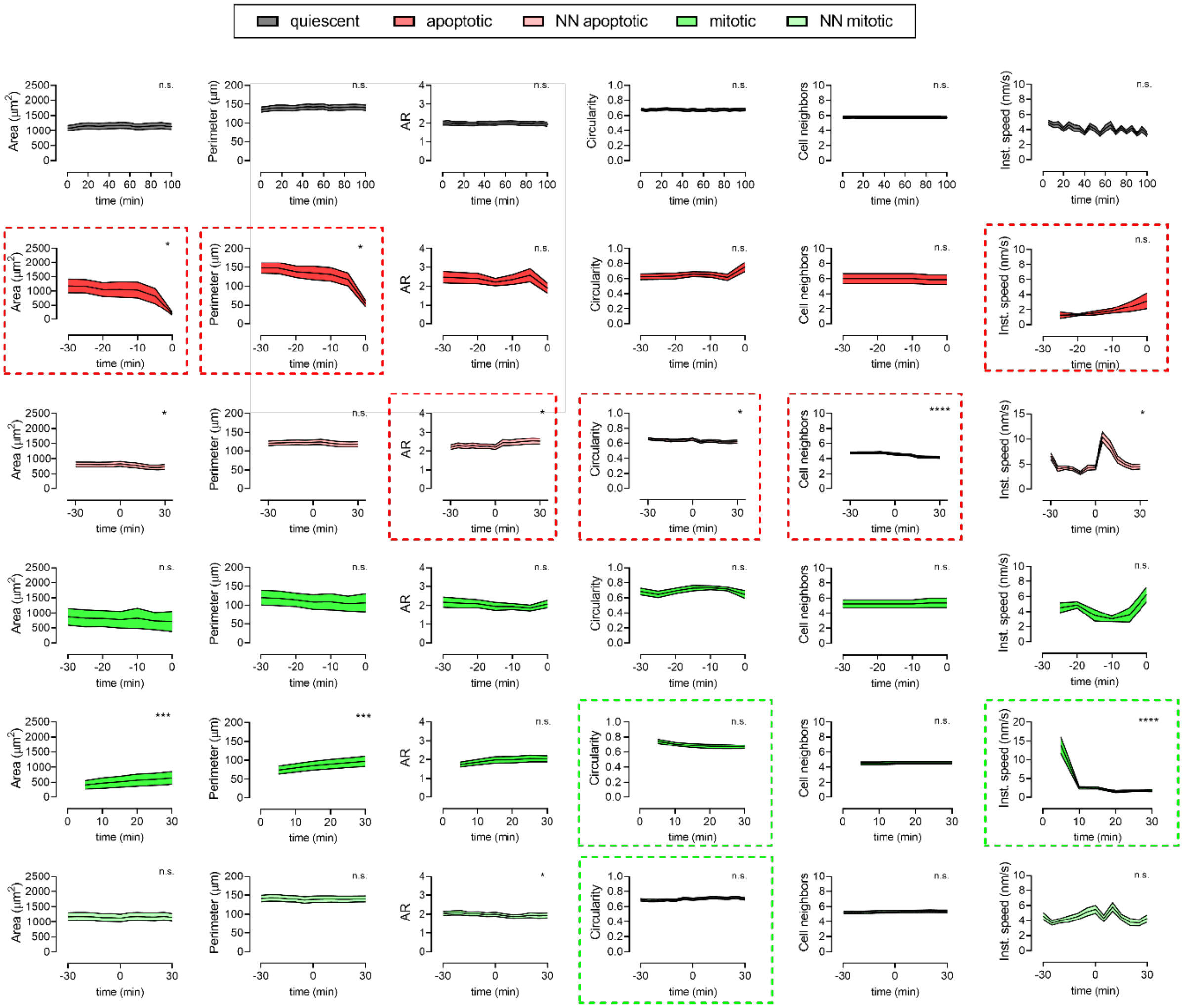


**Figure S9.** Time course of morphological metrics across quiescent, mitotic, and apoptotic cells (and their progeny and nearest neighbors) encompassing 183.5 hours of cellular imaging with 5-minute resolution. Data represent: quiescent cells ($n$ = 7) and their nearest neighbors ($n$ = 47) over 100 minutes, apoptotic cells ($n$ = 7) (over 30 minutes prior) and their nearest neighbors ($n$ = 39) (over 30 min before and after), mitotic cells ($n$ = 8) (over 30 minutes prior) and their paired daughter cells ($n$ = 8) (over 30 minutes after), and their nearest neighbors ($n$ = 43) (over 30 min before and after). Data are presented as mean ± SEM. Statistical analysis was performed using a Wilcoxon matched-pairs signed rank test to compare first and last time points (significant results denoted by asterisks) and using an F test to determine if linear regression displays a statistically non-zero slope (significant results denoted by dotted boxes): Data are presented as mean ± SEM. * $p < 0.05$, ** $p < 0.01$, *** $p < 0.001$, and **** $p < 0.0001$.

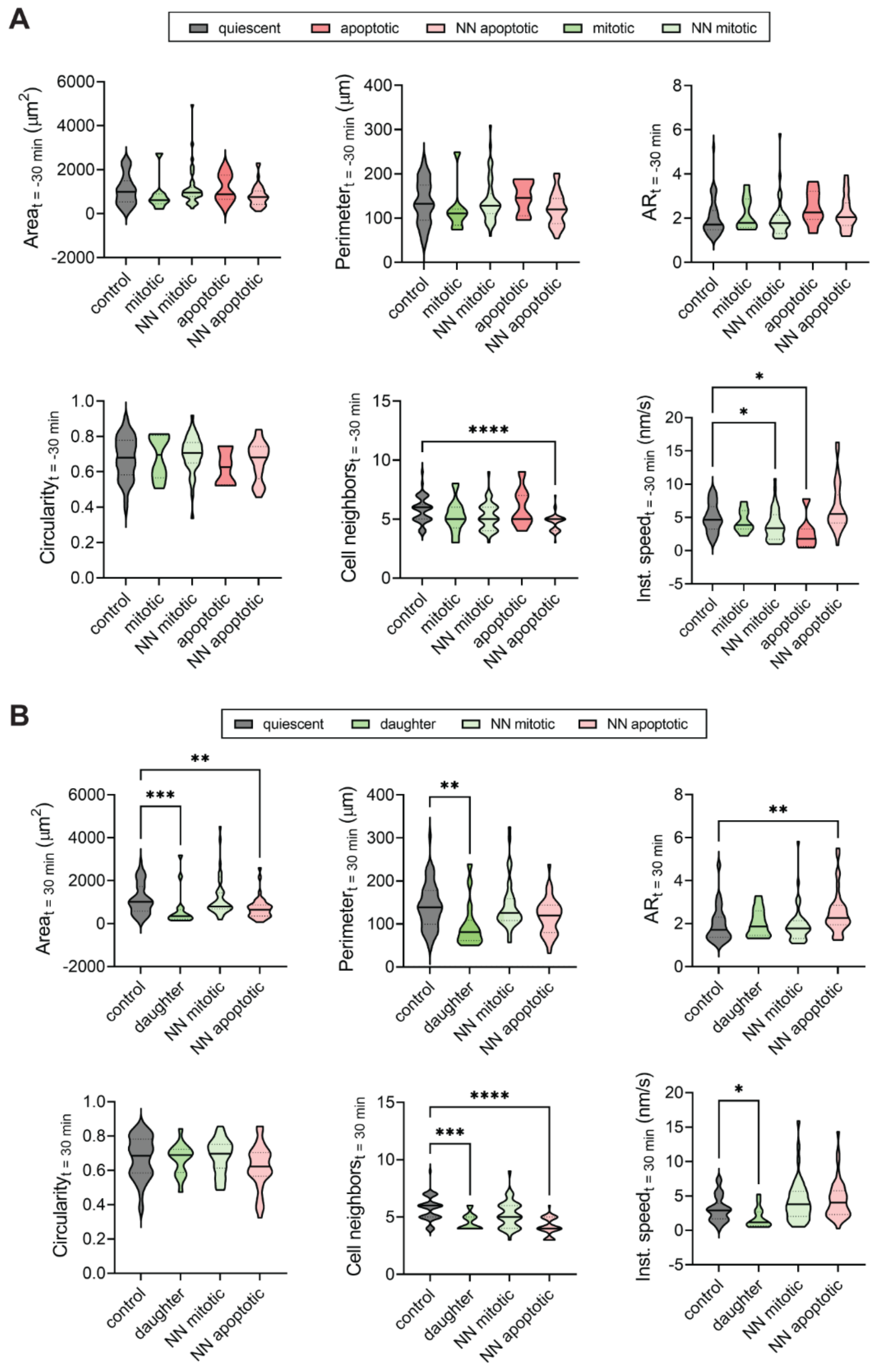


**Figure S10**. Violin plots of metrics across cell types at t = -30 minutes (A; top) and t = 30 minutes (B; bottom). Statistical analysis was performed using a Kruskal-Wallis test and post hoc Dunn's multiple comparisons test relative to control (quiescent cells): * $p < 0.05$, ** $p < 0.01$, *** $p < 0.001$, and **** $p < 0.0001$.

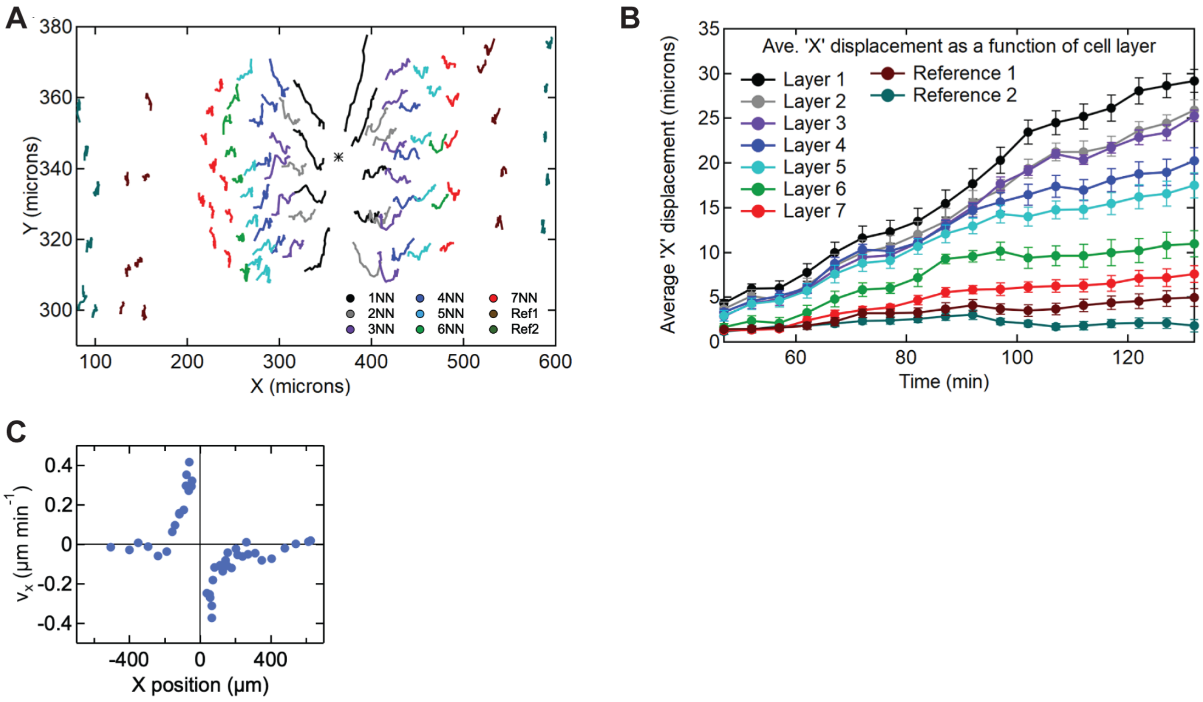


**Figure S11**. Response of three-dimensional iBMEC microvessels to wound formation by laser ablation.

(A) X and Y positions of cells at different near neighbor positions from the perimeter of the defect during wound closure. The trajectories of individual cells at different distances from the perimeter of the defect shows that cells within 6 - 7 near neighbors of the defect (about 200 $\mu$m) are involved in defect closing. Cells at the perimeter of the defect (first near neighbors, 1NN) show directed motion towards the point where the defect closes. Cells close to the perimeter (2NN to 7NN) show directed motion towards the defect, whereas cells further away (ref1 and ref 2) do not show direct motion. Black asterisk represents the centroid of the wound. Ref 1 denotes a reference point with cells 200 $\mu$m from defect center, and Ref 2 denotes a reference point with cells 250 $\mu$m from would center.

(B) X displacement versus time for cells at different distances from the wound. The x-component of the displacement (along the length of the microvessel) of individual cells towards the center of the defect increases approximately linearly with time up to closure. Data represent mean ± SE of cells tracked in S9A.

(C) The x-component of the cell velocity during defect closure was about 0.4 $\mu$m min$^{-1}$ for cells at the perimeter (1NN), but decreased with subsequent neighbors (2NN – 7NN). Cells more than 200 $\mu$m away from the defect show no global displacement during wound closure. The maximum speed is comparable to cell speeds reported for non-brain specific endothelial cells in conventional and microfluidic wound healing assays [1].

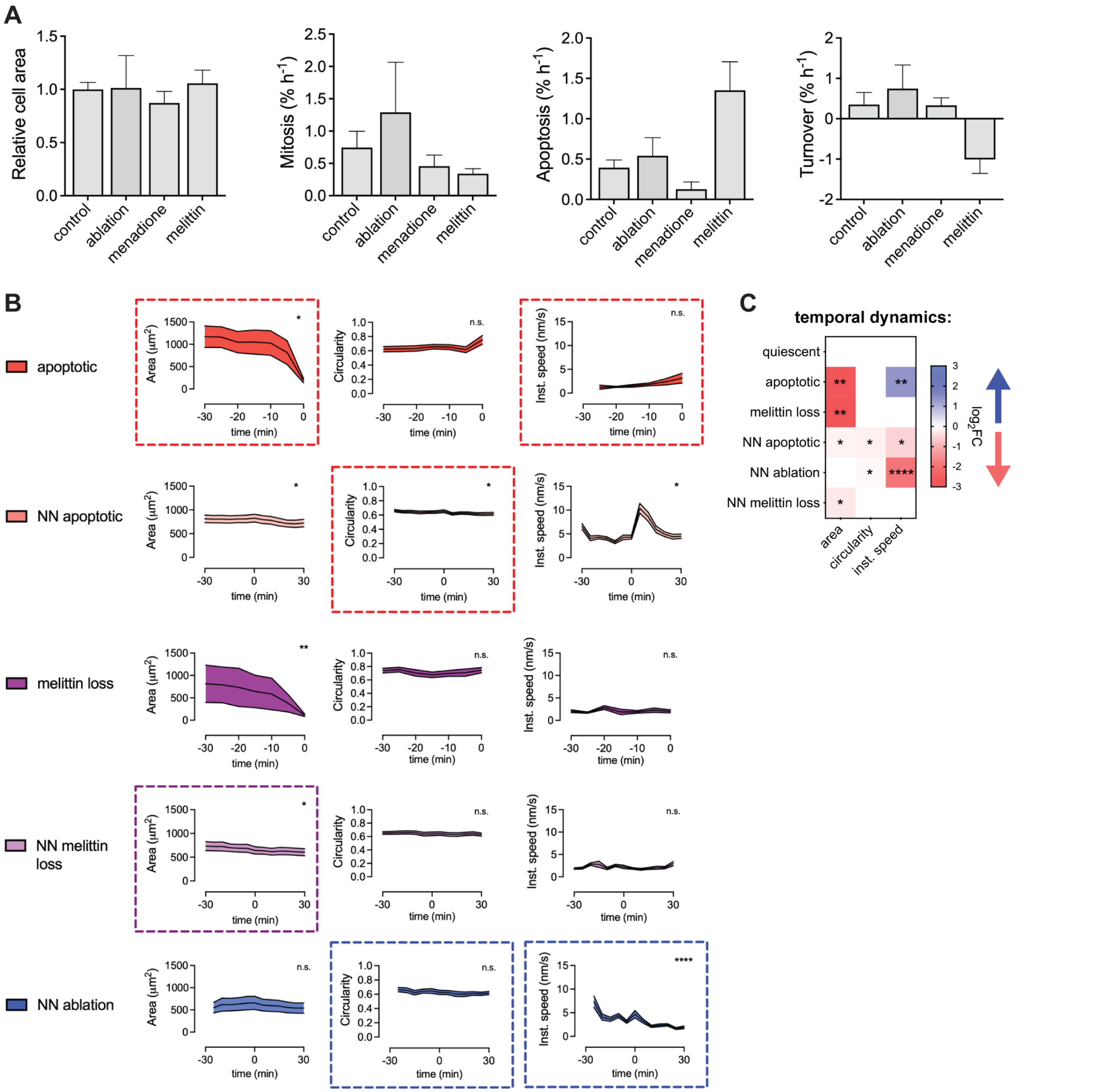


**Figure S12.** Comparison of cell turnover and morphology between homeostasis, ablation, menadione, and melittin exposure.

(A) Comparison of mitosis, apoptosis, and turnover rates where each condition displays similar starting average cell area. Data represents: control ($n$ = 7), ablation ($n$ = 3), menadione ($n$ = 4), and melittin ($n$ = 4), where $n$ represents distinct biological replicates (i.e. microvessels).

(B) Time course of area, circularity and instantaneous speed across conditions. Data represents: apoptotic cells ($n$ = 7) (over 30 minutes prior) and their nearest neighbors ($n$ = 39) (over 30 min before and after), cells lost during melittin exposure ($n$ = 8) (over 30 minutes prior) and their nearest neighbors ($n$ = 32) (over 30 min before and after), and nearest neighbors of a wound ($n$ = 18) (over 30 min before wound closure and 30 min after), where *n* represents individual cells across biological replicates. Statistical analysis was performed using a Wilcoxon matched-pairs signed rank test to compare first and last time points (significant results denoted by asterisks) and using an F test to determine if linear regression displays a statistically non-zero slope (significant results denoted by dotted boxes).

(C) Heatmap summarizing morphological changes across conditions. Significance and direction of change calculated using the maximally significant result between two statistical approaches mentioned above.

Data are presented as mean ± SEM. * $p < 0.05$, ** $p < 0.01$, and *** $p < 0.001$.

*Summary*. Turnover rates between microvessel conditions are similar, except following melittin exposure which causes a net loss of endothelial cells from the monolayer. Apoptotic cells show a decrease in area and increase in circularity immediately before apoptosis. The nearest neighbors of apoptotic cells show no change in morphology, but show an increase in instantaneous cell speed associated with migration into the area lost by the apoptotic cell. Cell loss in response to melittin exposure is characterized by a decrease in cell area prior to cell loss, but no change in circularity or instantaneous cell speed. The nearest neighbors of melittin-induced cell loss show no change in morphology or speed; healing of the defect occurs at longer time scales. Nearest neighbors of defects induced by laser ablation show no change in area or morphology, but show significant directed migration towards the centroid of the defect.

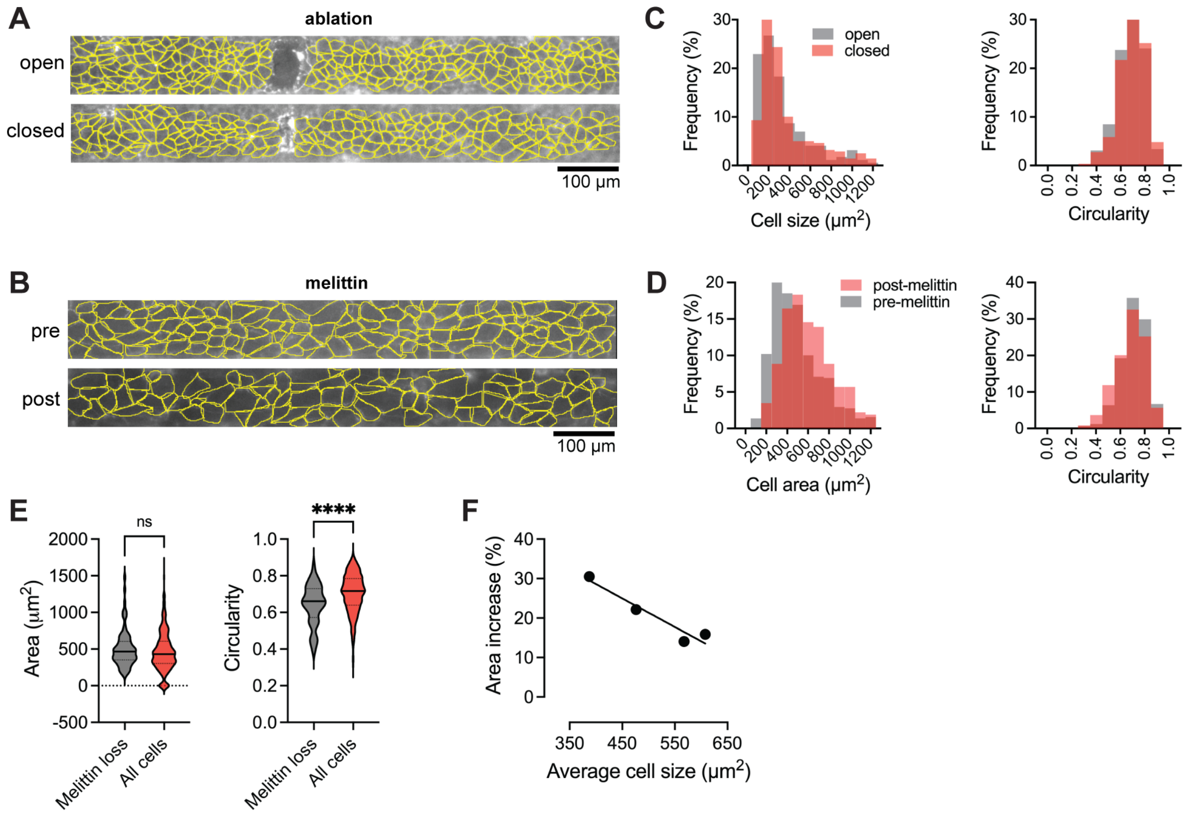


**Figure S13**. Monolayer area dynamics during wound healing and melittin exposure.

(A, C) Representative images showing traces of cells with a monolayer during an open versus closed wound and pre- and post-melittin exposure. Traces across replicates were used to generate histogram of cell size and circularity.

(B, D) Histograms of cell size and circularity following ablation and melittin exposure. $n$ = 353 cells in open wound and $n$ = 290 cells in closed wound; $n$ = 581 cells pre-melittin and 329 cells post-melittin.

(E) Initial size and circularity of cells lost during melittin exposure compared to the entire monolayer. $n$ = 75 cells lost during melittin exposure and $n$ = 581 cells pre-melittin.

(F) Changes in cell area in response to melittin relative to monolayer average cell size.

Significance calculated using a Mann-Whitney test. **** $p < 0.0001$.

**Table S1.** Summary of RNA sequencing meta-analysis.

| Ref. | Description | GEO # | Samples |
|---|---|---|---|
| [2] | iBMECs studied in a microfluidic model to probe the influence of shear stress | GSE129290 | GSM3704104, GSM3704105, GSM3704106, GSM3704107, GSM3704108, GSM3704109 |
| [3] | Human brain microvessels isolated by laser capture microdissection | GSE142209 | GSM4222739, GSM4222740, GSM4222741 |
| [4] | Immortalized BMECs (D3 line) | GSE76531 | GSM2027564, GSM2027565, GSM2027566, GSM2027567, GSM2027568, GSM2027569 |
| [5] | Immunopanned human BMECs | GSE73721 | GSM1901342, GSM1901343 |
| ENCODE Consortium [6] | Primary lung ECs | GSE78543 | GSM2072335, GSM2072334 |
| [7] | Primary human bronchial epithelial cells | GSE85402 | GSM2266295, GSM2266296, GSM2266297, GSM2266298, GSM2266299 |
| [8] | Caco-2 epithelial cell line | GSE164334 | GSM5006468, GSM5006469, GSM5006470 |
| [9] | iBMECs reprogrammed using ETS transcription factors | GSE138025 | GSM4096904 |
| [10] | iECs treated with DMSO or RepSox for 48 h | GSE142322 | GSM4225253, GSM4225254, GSM4225255, GSM4225256, GSM4225261, GSM4225262, GSM4225263, GSM4225264 |
| [11] | Endothelial progenitor cells (EPCs) treated with DMSO or Wnt agonist | GSE173206 | GSM5261814, GSM5261815, GSM5261816, GSM5261817, GSM5261818, GSM5261819, GSM5261820, GSM5261821 |

**Table S2.** Antibodies used in this study.

| Antibody | Vendor | Species | Cat. No | Dilution |
|---|---|---|---|---|
| Occludin | Invitrogen | Rabbit | 404700 | 1:100 |
| Claudin-5 * | Invitrogen | Mouse | 352500 | 1:200 |
| GLUT1 | Abcam | Rabbit | 115730 | 1:200 |
| P-gp | Sigma | Mouse | P7965 | 1:100 |
| CD31 | ThermoFisher | Rabbit | RB10333 | 1:25 |
| ICAM1 | Abcam | Mouse | AB2213 | 1:25 |
| VE-cadherin | R&D Systems | Goat | AF938 | 1:25 |
| CYP1A1 | Proteintech | Rabbit | 132411AP | 1:100 |
| MT2A | Abcam | Rabbit | AB192385 | 1:200 |
| CD271 | Miltenyi Biotec | Human (recombinant) | 130112790 | 1:50 |
| NG2 | ThermoFisher | Mouse | 53650482 | 1:20 |
| PDGFRβ | BD | Mouse | 558820 | 1:10 |

* Note: immunogen sequence used to generate claudin-5 antibody displays 100% sequence identity with claudin-5, while only 50% sequence identity with claudin-3, and no sequence identity with other claudin**s.**

### SUPPLEMENTAL DATA SETS

**Data S1.** Summary of bulk RNA-sequencing data. (Tab 1) Differentially expressed genes (DEGs) between iBMECs in 2D monolayers versus 3D microvessels. (Tab 2) DEGs between iBMECs cultured under 2.4 dyne $cm^{-2}$ shear stress versus static conditions. (Tab 3) DEGs between iBMECs in 3D microvessels exposed to TNFα for 24 hours versus controls. (Tab 4) Normalized enrichment scores (NES) for Molecular Signatures Database (MSigDB) hallmark gene sets for effects of 3D microenvironment and shear stress.

**Data S2**. Summary of tight junction dynamics data. (Tab 1) Principal component analysis of the morphological metrics that contribute to cell behavior. (Tab 2) Summary of time-dependent morphological metrics across cell categorizations (only statistically significant presented). (Tab 3) Summary of morphological metrics that differ across cell categorizations before and after cellular events (only statistically significant presented).

### Supplemental Notes:

*Note S1. Effect of media volume on iBMEC phenotype*

Motivated by a desire to minimize reagent use while maintaining cellular fidelity, we differentiated iBMECs in parallel using 1 mL or 2 mL of medium to determine impact of media volume on cell phenotype (**Fig. S1B)**. Previous studies have reported the influence of numerous variables on iBMEC differentiation, including hypoxia [29], intermediate mesodermal induction [21, 32], media composition [33], serum free alternatives [34], source/choice of ECM coating material [35-37], and iPSC seeding density [38]. Reduced media volume increased the fraction of all singularized cells which attached to sub-culture plates (p = 0.023), suggesting increased purity of endothelial-like cells resulting from the differentiation (**Fig. S1C**). To determine functional differences between monolayers generated using different media volumes, iBMECs were seeded onto 2D transwells to determine the transendothelial electrical resistance (TEER) and permeability of Lucifer yellow (444 Da), 10 kDa dextran, glucose, and rhodamine 123 (R123). Differentiation in 1 mL volume resulted in monolayers with significantly increased TEER over six days of culture (Fig. S1D). The average TEER was increased for each cell line ($p < 0.001$, $p < 0.001$, $p = 0.001$, for TJ, PM and ACTB, respectively). The average difference in TEER was ~2-fold, with similar differences observed across all cell lines and across all time points (**Fig. S1E**).

Consistent with TEER results, Lucifer yellow permeability was lower in monolayers of iBMECs differentiated using 1 mL volume at day 2 ($p = 0.043$) (**Fig. S1F**). After six days of culture on transwells, permeability increased ~6-fold for both media volumes ($p$ = 0.056 and 0.078, respectively), and permeability was not significantly different between the two media volumes ($p = 0.450$). 10 kDa dextran permeability displayed similar trends, however, no substantial differences were observed between media volumes at either day 2 or day 6 (**Fig. S1G**). These results suggest that paracellular barrier strength is improved for only small compounds using 1 mL media volume, and that for both media volumes paracellular barrier strength decreases in less than one week of transwell culture in the absence of supporting cells.

We next assessed the influence of media volume on two transcellular transport pathways: glucose transport and P-gp efflux (**Fig. S1H**). The GLUT-1 transporter (*SLC2A1*) is responsible for transporting glucose into brain parenchyma. We observed that glucose transport preferentially occurred in the apical-to-basolateral direction for both media volumes suggesting physiological localization and function of GLUT1. However, despite an was ~2-fold higher directional transport for monolayers of 1-mL iBMECs the difference was not statistically significant ($p$ = 0.353). P-gp (*ABCB1*) is an efflux pump for which Rhodamine 123 is a substrate. We observed that R123 transport preferentially occurred in the basolateral-to-apical direction for both media volumes and was higher for monolayers of 1-mL iBMECs but was not statistically significant ($p$ = 0.054).

iBMECs across both media volumes displayed similar localization of endothelial markers (CD31), tight junction proteins (ZO1, Occludin, Claudin-5), nutrient transporters (GLUT1), and efflux transporters (P-gp) (**Fig. S1I**). Using a semi-quantitative analysis of relative protein expression (average fluorescence intensity), we did not observed differences across key BBB markers, beyond a slight reduction in GLUT1 signal for 1 mL-iBMECs (**Fig. S1J**). RNA sequencing was utilized to compare global gene expression of source iPSCs and iBMECs generated using 1- or 2-mL media volume. As previously reported [12], differentiation of iBMECs is associated with widespread changes in gene expression including upregulation of BBB markers (e.g. *SLC2A1*) and endothelial cell markers (e.g. *CDH5*), and downregulation of pluripotency markers (e.g. *POU5F1*, *SOX2*, and *MYC*). 1 mL and 2 mL iBMECs display similar gene expression profiles with only 59 DEGs (**Fig. S1K**).

*Note S2. Variability of iBMEC differentiation*

Variability in iPSC differentiation is a major limitation for reproducible disease modeling [13, 14]. For example, serum lot and iPSC source have been shown to introduce variability into iBMEC differentiation outcomes [15, 16], suggesting that serum-free alternatives and optimization studies are needed to achieve high levels of reproducibility in independent laboratories. Additionally, cryopreservation strategies that maintain iBMEC phenotype can be used to reduce batch-to-batch variability [12, 17]. To confirm the robustness of our findings, we compared TEER of 1 mL and 2 mL iBMECs after cryopreservation, using an independent serum lot, and using an independent iPSC source. TEER values were higher using 1 mL media volume for all variables (**Fig. S2A**). Following published cryopreservation approaches [12], similar peak TEER values were achieved (2,540 ± 380 Ω $cm^2$). To probe batch-to-batch variability of iBMEC differentiations we constructed Bland Altman plots of average TEER for monolayers of 1 mL iBMECs versus difference in average TEER between media volumes. Across 55 independent differentiations, 91% of differentiations produced higher TEER values for iBMEC monolayers using 1 mL media volume, and the difference was most pronounced when TEER values for 1 mL iBMEC were highest (**Fig. S2B**). Across our extensive paracellular permeability dataset, we further probed the relationship between TEER and permeability, finding that Lucifer yellow permeability was nearly constant (~2 x $10^{-6}$ cm $s^{-1}$) above TEER values of ~500 Ω $cm^2$, while the permeability of 10 kDa dextran was nearly constant (~5 x $10^{-8}$ cm $s^{-1}$) above TEER of ~250 Ω $cm^2$ (**Fig. S2C**). Below these cut-offs, the permeability increased sharply with decreasing TEER. The permeabilities of Lucifer yellow and 10 kDa dextran are linearly related across a wide range of permeabilities ($r^2$ = 0.97), where 10 kDa dextran has ~50-fold lower permeability (Lucifer yellow has ~20-fold lower molecular weight) (**Fig. S2D**). Collectively, these results provide quantitative metrics to define the reproducibility of iBMECs, for example, a threshold TEER value can be used to reduce variability when screening drug permeability.

*Note S3. Mechanisms of media volume effect on iBMEC phenotype*

As media volumes were altered during both the UM/F- and EC+RA phase of the differentiation, we first sought to determine if the effect of media volume was predominately due to one media phase. When media volume was toggled between 1 mL and 2 mL for the UM/F- (6 days) and EC+RA (2 days) phases, we found that the UM/F- phase solely accounted for improved iBMEC phenotype. TEER time course was similar regardless of EC+RA media volume, while UM/F- media volume produced an ~2-fold difference (**Fig. S3A**). Additionally, the average TEER was irrespective of EC+RA media volume ($p$ = 0.999 and 0.997, for 1 mL and 2 mL UM/F- respectively) (**Fig. S3B**).

We hypothesized that media volume may alter nutrient levels and oxygenation and within the cell culture media. Glucose levels were recorded at the first and last day of UM/F- differentiation from conditioned differentiation media, and across switch frequencies. All recordings were ~6 mM for spent media with ~16 mM for fresh media, suggesting that difference in glucose are not responsible for iBMEC phenotypic improvements (**Fig. S3C**). Additionally, oxygen levels were recorded at the first and last day of UM/F- differentiation, however, there was no difference between media volumes ($p$ = 0.811 and 0.448, at day 2 and 6 respectively) (**Fig. S3D**). Oxygenation slightly decreased overtime for both media volumes but was roughly similar to atmospheric levels (~21%) (**Fig. S3E**).

To further probe the effects of media volume on iBMEC phenotype, we also altered the frequency of media switches. While under baseline conditions media switches are conducted daily during UM/F-, we found that frequency of media switches modulated TEER improvements (**Fig. S3F**). Increasing media switch frequency during the 1 mL differentiation abolished TEER improvements ($p$ = 0.019), while reducing media switch frequency during the 2 mL differentiation increased TEER ($p$ = 0.020). Thus, both reduction of media volume or reductions in frequency of media switches improves iBMEC phenotype, while minimizing media consumption (i.e. material costs).

*Note S4. Calculation of permeability in 2D and 3D.*

The permeability associated with diffuse transport of a solute across a membrane, such as a cell monolayer, is obtained from Fick's first law [18-23]. The method for calculating permeability of a fluorescent solute in a 3D microvessel is similar, and has been described in detail elsewhere [24-26]. In both cases, the permeability is given by:

$$P = \frac{1}{n_0}\left(\frac{dn_{out}}{dt}\right)\frac{V}{A}$$

where $n_0$ is the number of molecules on the input side (apical chamber or lumen), $V$ is the volume on the input side (apical chamber or lumen), $dn_{out}/dt$ is the change in concentration on the output side (basolateral chamber or ECM), and $A$ is the area (membrane or vessel lumen). The derivation assumes that: (1) the concentration of solute in the input chamber ($n_0/V$) is approximately constant, (2) transport from the output side to the input side can be neglected ($c_{in} > c_{out}$; usually equivalent to short times), and (3) transport is dominated by passive diffusion across the cell monolayer. In comparing values of permeability there are three experimental factors that could lead to differences in values between 2D and 3D measurements: (1) laminar flow/shear stress, (2) cell-ECM interactions, and (3) curvature. Lastly, in 2D Transwells, poor adhesion of the monolayer to the walls of the Transwell support can lead to a short circuit path or perimeter effect [18]. The continuous boundary conditions around the lumen of a 3D microvessel eliminate this possible source of error.

**Supplemental Methods**:

*iBMEC differentiation*

iPSCs were plated at 10,000 cells $cm^{-2}$ on Matrigel-coated six-well plates and grown for two days in mTESR1 (StemCell Technologies) with 10 μM ROCK inhibitor Y27632 (RI; ATCC) supplemented for the initial 24 hours. Subsequently, cells were treated with unconditioned media without bFGF (UM/F-): DMEM/F12 (Life Technologies) supplemented with 20% knockout serum replacement (Life Technologies), 1% non-essential amino acids (Life Technologies), 0.5% GlutaMAX (Life Technologies) and 0.836 μM beta-mercaptoethanol (Life Technologies) for six days, and retinoic acid-supplemented endothelial media (EC+RA): human endothelial cell serum-free media (Life Technologies) supplemented with 1% human platelet poor derived serum (Sigma), 2 ng $mL^{-1}$ bFGF (R&D Systems), and 10 μM all-trans retinoic acid (Sigma) for two days. 2 mL volume was used for daily mTESR1 media switches; 1 mL or 2 mL volumes were used for daily UM/F- media switches; 1 mL or 2 mL volumes were used for EC+RA media, with no media switch over the two-day EC+RA treatment period. Post-differentiation iBMECs were sub-cultured onto 6-well tissue culture plates at 1:1 surface area ratio using EC+RA media supplemented with 1% pen-strep and 10 μM ROCK inhibitor Y27632. After one-hour media was switched to remove non-adherent cells. One day after initial subculture, media was switched to basal media. A PowerPlex 18D kit (Promega) was used to determine the short tandem repeat (STR) profiles of cell lines; all lines were confirmed to be isogenic to familial WTC iPSCs. A PCR based MycoDtect kit (Greiner Bio-One) was used to confirm cell lines were not contaminated with mycoplasma.

*Transwell barrier characterization*

After differentiation, cells were singularized using Accutase and sub-cultured onto surfaces treated overnight with 50 μg $mL^{-1}$ human placental collagen IV (Sigma) and 25 μg $mL^{-1}$ fibronectin from human plasma (Sigma). Cells were seeded at a density of 1 x $10^6$ $cm^{-2}$ on transwells prior to transendothelial electrical resistance (TEER) recordings using an EndOhm (World Precision Instruments), as previously reported [16]. All measurements were conducted on 6.5 mm transwells with a 0.4 μm pore polyester membrane insert immediately after removal from an incubator, were corrected against values of an empty transwell insert, and normalized by transwell surface area.

Lucifer yellow, 10 kDa dextran, Rhodamine 123 and glucose permeability was measured on day 2; Lucifer yellow and 10 kDa dextran was also measured on day 6. 200 μM Lucifer yellow (LY;

ThermoFisher), 2 µM 10 kDa dextran (ThermoFisher), 10 µM rhodamine 123 (R123; ThermoFisher), and 25 mM d-glucose (Sigma) were prepared in transport buffer comprised of distilled water with 0.12 M NaCl, 25 mM $NaHCO_3$, 2 mM $CaCl_2$, 3 mM KCl, 0.4 mM $K_2HPO_4$, 2 mM $MgSO_4$, and 1 mM HEPES supplemented with 0.1% human platelet poor derived serum. For some experiments, 10 kDa dextran was multiplexed with Lucifer yellow or Rhodamine 123, which have non-overlapping fluorescent spectra. To measure apical to basolateral permeability, transport buffer containing the fluorophore of interest was added to the apical chamber (100 µL), while the basolateral chamber contained 600 µL of transport buffer. To measure the basolateral-to-apical permeability, transport buffer containing the fluorophore was added to the basolateral chamber (600 µL), while the apical chamber contained 100 µL of transport buffer. For Lucifer yellow and 10 kDa dextran, apical-to-basolateral permeability was measured at 15 and 30 minutes. For rhodamine 123 and 10 kDa dextran, both apical-to-basolateral and basolateral-to-apical permeability was measured at 60 and 90 minutes. For glucose, both apical-to-basolateral and basolateral-to-apical permeability was measured at 5, 10, 15, and 30 minutes. Time courses were selected within the linear regime of transport [18], and conducted under gentle agitation at 37 °C. For Lucifer yellow, 10kDa dextran and rhodamine 123, transport buffer was collected at each time point and fluorescence measured using a Synergy™ H4 microplate reader (Biotek) with the following excitation and emission settings 428 nm / 545 nm, 647 nm / 667 nm, and 495 nm / 525, respectively. For glucose, transport buffer was collected and quantified using a glucose colorimetric detection kit (ThermoFisher Scientific), utilizing absorbance recordings at 560 nm. Concentration was determined from calibration curves based on serial dilutions spanning four orders of magnitude. The apparent permeability of each probe was calculated as previously reported [27, 28].

*Oxygen and glucose recordings*

Dissolved oxygen (DO) in the pericellular region at the base of the well was measured continuously by an OXY-4 mini meter (PreSens) capable of optically reading DO levels from a sensor spot patch placed in a six-well plate. Media volumes (1 mL and 2 mL) were evaluated simultaneously. Measurements were performed within an incubator at 37°C and 5% $CO_2$ over a 24 hour period following media switches on the first and last day of the UM/F- differentiation phase. To quantify the concentration of D-glucose in the conditioned media, 1 mL of media was collected from wells of iBMECs undergoing differentiation under each of the media volume conditions described above, on both days 0 and 6 of the differentiation (at each media transition stage). All samples were immediately frozen at -20 °C and thawed for 2 hours at room temperature prior to quantification using a glucose colorimetric detection kit (ThermoFisher Scientific) as previously described.

*Pericyte differentiation and characterization*

Brain pericyte-like cells (iPCs) were derived through a neural crest intermediate with minor modification to published protocols [29]. Briefly, WTC-iPSC-PMs were singularized with Accutase and seeded at 9.1 × $10^4$ cells $cm^{-2}$ onto Matrigel-coated plates with mTESR1 + 10 $\mu$M Y27632. Differentiation to neural crest stem cells (NCSC) began the following day (day 0) in E6 media (StemCell Technologies) supplemented with 22.5 mg $L^{-1}$ heparin sodium salt from porcine mucosa (Thermo Fisher Scientific), 10 $\mu$g $L^{-1}$ bFGF (Fisher Scientific), 1 $\mu$M CHIR99021 (Peprotech), 10 $\mu$M SB431542 (Peprotech), and 1 $\mu$M dorsomorphin (Peprotech) (termed E6-CSFD). Media was replaced daily, and once reaching confluence (2-3 days initially, then approximately every 4 days), cells were dissociated with Accutase, split 1:12, and reseeded on Matrigel-coated plates. On day 15, NCSCs were enriched by magnetic-activated cell sorting (MACS). Briefly, cells were dissociated with Accutase, passed through a 40 $\mu$m strainer, and labeled using anti-CD271 (LNGFR)-PE (Miltenyi Biotec 130-112-790) in MACS buffer (0.5% bovine serum albumin + 2 mM EDTA in sterile PBS) at 4°C for 15 minutes. Cells were washed with MACS buffer, and then incubated with anti-PE microbeads (Miltenyi Biotec 130-048-801) in MACS buffer at 4°C for 15 min. Cells were again washed, then run through an MS column held by a Mini-MACS Separator (Miltenyi Biotec). CD271+ NCSCs were seeded at 1 × $10^4$ cells $cm^{-2}$ on uncoated six-well plates in E6-CSFD + 10 $\mu$M Y27632. Differentiation towards a pericyte lineage was initiated the next day (day 16) by switching to E6 media supplemented with 10% FBS. Media was replaced daily until day 25, during which cells were routinely passaged 1:2 once reaching confluence (without Y27632 supplementation). For flow cytometric evaluation, cells were incubated with Alexa Flour-488 conjugated NG2 and PE-conjugated PDGFRβ antibodies in MACS buffer on ice for 30 min and then washed three times. Marker expression was measured by a BD FACS Canto flow cytometer, were forward-side scatter plots were used to exclude dead cells and all analyses were performed against corresponding isotype controls.

[15] E.H. Neal, N.A. Marinelli, Y. Shi, P.M. McClatchey, K.M. Balotin, D.R. Gullett, K.A. Hagerla, A.B. Bowman, K.C. Ess, J.P. Wikswo, E.S. Lippmann, A Simplified, Fully Defined Differentiation Scheme for

Producing Blood-Brain Barrier Endothelial Cells from Human iPSCs, Stem Cell Reports 12(6) (2019) 1380-1388.
[16] G.N. Grifno, A.M. Farrell, R.M. Linville, D. Arevalo, J.H. Kim, L. Gu, P.C. Searson, Tissue-engineered blood-brain barrier models via directed differentiation of human induced pluripotent stem cells, Sci Rep 9(1) (2019) 13957.
[17] H.K. Wilson, M.G. Faubion, M.K. Hjortness, S.P. Palecek, E.V. Shusta, Cryopreservation of Brain Endothelial Cells Derived from Human Induced Pluripotent Stem Cells Is Enhanced by Rho-Associated Coiled Coil-Containing Kinase Inhibition, Tissue Eng Part C Methods 22(12) (2016) 1085-1094.
[18] A.D. Wong, M. Ye, A.F. Levy, J.D. Rothstein, D.E. Bergles, P.C. Searson, The blood-brain barrier: an engineering perspective, Frontiers in neuroengineering 6 (2013) 7.
[19] T.T. Tran, A. Mittal, T. Gales, B. Maleeff, T. Aldinger, J.W. Polli, A. Ayrton, H. Ellens, J. Bentz, Exact kinetic analysis of passive transport across a polarized confluent MDCK cell monolayer modeled as a single barrier, J Pharm Sci 93(8) (2004) 2108-23.
[20] A. Siflinger-Birnboim, P.J. Del Vecchio, J.A. Cooper, F.A. Blumenstock, J.M. Shepard, A.B. Malik, Molecular sieving characteristics of the cultured endothelial monolayer, J Cell Physiol 132(1) (1987) 111-7.
[21] J. Karlsson, P. Artursson, A Method for the Determination of Cellular Permeability Coefficients and Aqueous Boundary-Layer Thickness in Monolayers of Intestinal Epithelial (Caco-2) Cells Grown in Permeable Filter Chambers, Int J Pharm 71(1-2) (1991) 55-64.
[22] P.H. Barry, J.M. Diamond, Effects of unstirred layers on membrane phenomena, Physiol Rev 64(3) (1984) 763-872.
[23] A. Avdeef, Physicochemical profiling (solubility, permeability and charge state), Curr Top Med Chem 1(4) (2001) 277-351.
[24] F.E. Curry, V.H. Huxley, R.H. Adamson, Permeability of single capillaries to intermediate-sized colored solutes, Am J Physiol 245(3) (1983) H495-505.
[25] V.H. Huxley, F.E. Curry, R.H. Adamson, Quantitative fluorescence microscopy on single capillaries: alpha-lactalbumin transport, Am J Physiol 252(1 Pt 2) (1987) H188-H197.
[26] Y. Yuan, W.M. Chilian, H.J. Grainger, D.C. Zawieja, Permeabiility to albumin in isolated coranary venules, American Journal of Physiology 264(34) (1993) 543-552.
[27] M.E. Katt, Z.S. Xu, S. Gerecht, P.C. Searson, Human Brain Microvascular Endothelial Cells Derived from the BC1 iPS Cell Line Exhibit a Blood-Brain Barrier Phenotype, PLoS One 11(4) (2016) e0152105.
[28] M.E. Katt, L.N. Mayo, S.E. Ellis, V. Mahairaki, J.D. Rothstein, L. Cheng, P.C. Searson, The role of mutations associated with familial neurodegenerative disorders on blood-brain barrier function in an iPSC model, Fluids Barriers CNS 16(1) (2019) 20.
[29] M.J. Stebbins, B.D. Gastfriend, S.G. Canfield, M.S. Lee, D. Richards, M.G. Faubion, W.J. Li, R. Daneman, S.P. Palecek, E.V. Shusta, Human pluripotent stem cell-derived brain pericyte-like cells induce blood-brain barrier properties, Sci Adv 5(3) (2019) eaau7375.
